# Theoretical Analysis of Principal Components in an Umbrella Model of Intraspecific Evolution

**DOI:** 10.1101/2021.11.28.470252

**Authors:** Maxime Estavoyer, Olivier François

## Abstract

Principal component analysis (PCA) is one of the most frequently-used approach to describe population structure from multilocus genotype data. Regarding geographic range expansions of modern humans, interpretations of PCA have, however, been questioned, as there is uncertainty about the wave-like patterns that have been observed in principal components. It has indeed been argued that wave-like patterns are mathematical artifacts that arise generally when PCA is applied to data in which genetic differentiation increases with geographic distance. Here, we present an alternative theory for the observation of wave-like patterns in PCA. We study a coalescent model – the umbrella model – for the diffusion of genetic variants. The model is based on genetic drift without any particular geographical structure. In the umbrella model, splits from an ancestral population occur almost continuously in time, giving birth to small daughter populations at a regular pace. Our results provide detailed mathematical descriptions of eigenvalues and eigenvectors for the PCA of sampled genomic sequences under the model. Removing variants uniquely represented in the sample, the PCA eigenvectors are defined as cosine functions of increasing periodicity, reproducing wave-like patterns observed in equilibrium isolation-by-distance models. Including rare variants in the analysis, the eigenvectors corresponding to the largest eigenvalues exhibit complex wave shapes. The accuracy of our predictions is further investigated with coalescent simulations. Our analysis supports the hypothesis that highly structured wave-like patterns could arise from genetic drift only, and may not always be artificial outcomes of spatially structured data. Genomic data related to the peopling of the Americas are reanalyzed in the light of our new theory.

## Introduction

Investigating population structure from large genotypic data set is one of the most important steps of modern population genetic data analysis. This step is often carried out with principal component analysis (PCA). For a sample of size n, PCA analyzes population genetic structure by computing the eigenvalues and eigenvectors of the empirical covariance matrix [23, 26]. The projection of samples on eigenvectors are called principal components (PCs). The first PCs represent the axes of genetic variation which carry the most information, and the eigenvalues represent the variances of samples along the axes. The use of PCA dates back to the early days of human population genetics [5, 35], and it has been shown theoretically that it can reveal information about admixture and population differentiation among samples [37, 34, 38, 18].

Because PCA is an exploratory method that makes no assumptions about population history, interpretation of results in terms of demographic events is sometimes difficult, in particular when the details of a range expansion are scrutinized [36, 34, 16, 1,3]. More precisely, some authors interpreted gradient and wave patterns in these PCs as signatures of specific migration events [35]. But Novembre and Stephens reported that highly structured wave-like patterns that are seen in PCA when genetic data are simulated under range expansion models, are also observed under equilibrium isolation-by-distance models without range expansions [36]. Their study warned that the wave-like patterns could be mathematical artifacts that arise when PCA is applied to spatial data in which covariance between locations decays with geographic distance [10, 19]. Sinusoidal shapes similar to those observed with geographic samples are also observed with temporal samples of ancient DNA genotypes [17], and the combination of time and spatial heterogeneity in sampling further complicates the interpretation of patterns observed in PC plots [11]. Those difficulties have questioned the usefulness of PCA for species that evolved under geographic models of range expansion for which wave-like patterns were first observed.

Range expansions are, however, common in the evolution of species. For example, they include the worldwide expansion of monarch butterflies [40], or the invasion of Cane toads in Australia [39]. The models have been extensively studied theoretically and with computer simulations [2, 22, 13]. Humans are one of the most studied example of range expansion, exhibiting a positive correlation between genetic and geographic distance at the worldwide scale [42], and a proposed model for that pattern involved serial founder effects [9, 6]. In the serial founder effect model, populations migrate outward from a geographic origin through a process where each of a series of populations is formed from a subset of the previous population in the expansion [6]. Evolution under the model corresponds to a population tree having maximal imbalance (Figure 1), and several patterns of human genetic variation were explained by serial colonization out of Africa [9, 6, 7, 25].

**Figure 1.**
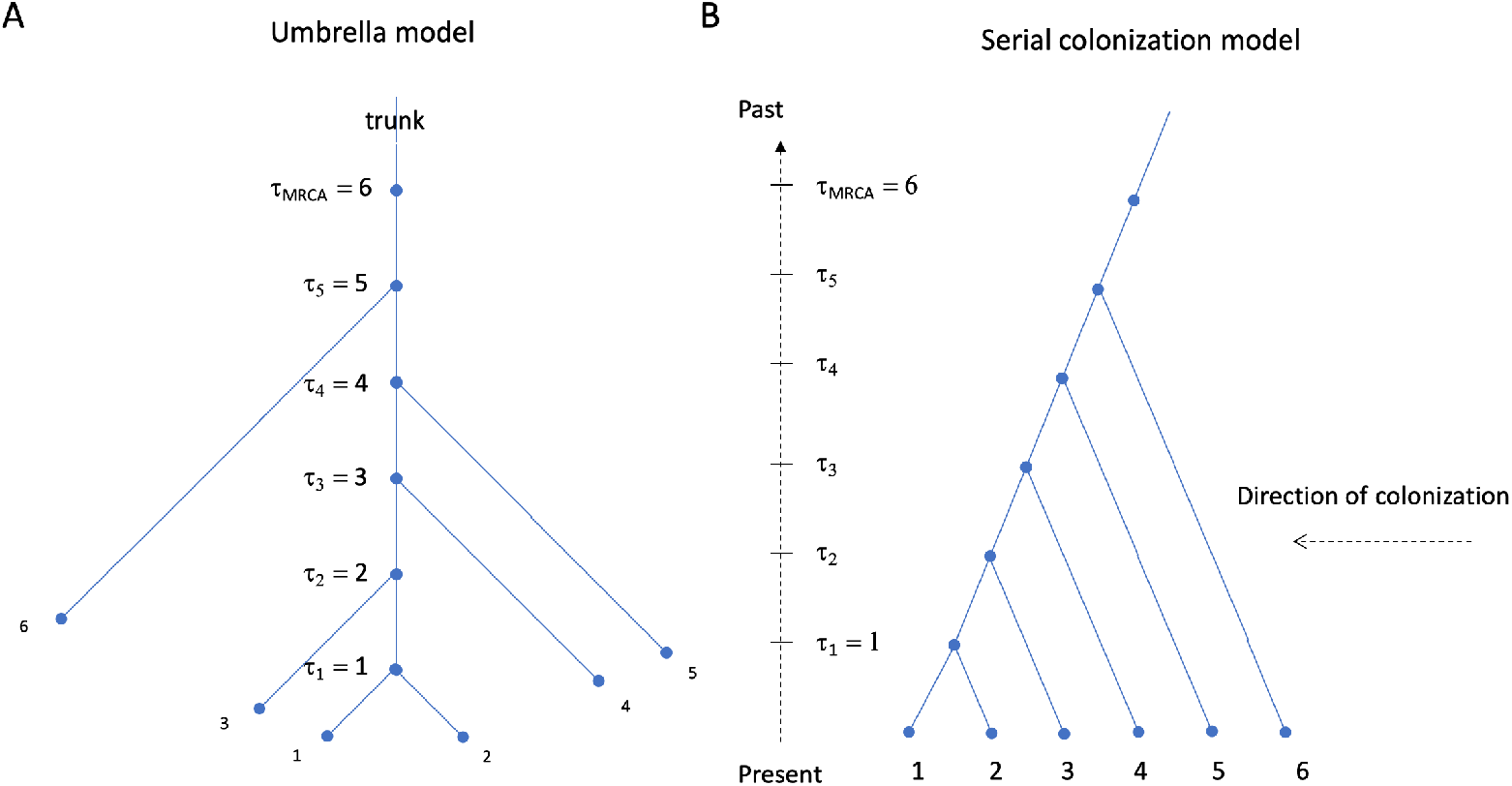
Umbrella Model. (**A**) Skeleton tree for the umbrella model: The individuals at the leaves are labelled according to the time of divergence with the ancestral lineage. Individual lineages coalesce with the ancestral lineage at their time of divergence plus one unit. (**B**) Equivalent description of the UM as a serial colonization model, in which individuals at the leaves are labelled according to their position in a linearly arranged habitat. The initial population is given the highest label. The value *τ_k_* indicates the splitting time for individual lineage *k* +1 (*τ*_1_ is for lineages 1 and 2).

In this study, we analyze a simple model of genetic drift in which small populations diverge from an ancestral population at a constant pace, through a process similar to a serial colonization model. The model is called the *umbrella model* in reference to a tree known as the umbrella pine, parasol pine or Italian stone pine, in which the length of a branch decreases with the distance to the root. Unlike the serial founder effect model, the samples are not necessarily localized in geographic space, and patterns of genetic variation result from temporal drift only [43]. The model describes the diffusion of genetic variants based on hierarchy of fissions from a genetically homogeneous population without any geographical structure in the sample. The samples sort according to their date of coalescence within the trunk of the tree, representing the ancestral population lineage. Removing variants that occur exactly once in the sample, the model also describes the neutral evolution of a genetically homogeneous population since the most recent common ancestor of the sample. In this case, the present-day sequences are not different of ancient DNA sequences sampled at their date of coalescence with the ancestral lineage. Thus the model is not only relevant to present-day population structure analysis, but also highlights some interesting properties of population structure analysis for ancient DNA samples.

We provide a detailed mathematical description of population structure under the umbrella model by studying the PCA eigenvalues and eigenvectors of sampled genomic sequences. The eigenvalues and eigenvectors of the PCA will be obtained after approximating the finite-size empirical covariance matrix by an integral (Hilbertian) operator. Removing variants that are uniquely represented in the sample, the eigenvectors are defined as cosine functions of increasing periodicity, reproducing wave-like patterns observed in equilibrium isolation-by-distance models. Keeping those rare variants produces complex oscillatory eigenvectors, for which mathematical descriptions are obtained for the largest eigenvalues. The accuracy of approximation will be further investigated with coalescent simulations, and genomic data related to the settlement of the Americas will be reanalyzed in the light of the new results [33].

## 1 Model description and theoretical background

### 1.1 The umbrella model

We study the genealogical relationships of *n* genomic sequences obtained from haploid individuals under a serial colonization model of evolution, the umbrella model (UM).

#### Skeleton tree

The UM defines a skeleton tree for the coalescent genealogies of the *n* samples. The skeleton can be viewed as an approximation of an ancestral population that splits continuously in time. The model considers *n* populations of equal size *N*, which must be thought of as being small compared to the effective population size *N_e_* (*N* is of order *N_e_/n*). Time is measured in units of population size. Temporal evolution is defined by population splits that occur each unit of time. Example values related to human data may be around *N_e_* = 10, 000, *N* = 100 and *n* =100. A splitting event gives birth to an external branch that diverges from the trunk of the tree (Figure 1A). The geographic localization of the daughter population can be arbitrary, and no relationship between geographic and genetic distance is modelled a priori. The skeleton tree of the UM has *n* external branches, for which the total length is equal to 1 + *n*(*n* – 1)/2, and *n* — 1 internal branches of unit length, forming a trunk of length *n*. The total tree length is equal to *n*(*n* + 1)/2.

#### Sample genealogy

In the UM, a single individual is sampled from each population, so that the sample size, n, is equal to the number of populations. The average time since the most recent common ancestor of all individuals is equal to *n*. Individual lineages coalesce with the ancestral lineage at their time of divergence plus one unit. Thus the samples sort according to their date of coalescence within the trunk of the tree. More precisely, individual lineage *i* (*i* ≥ 2) coalesces with the ancestral lineage at time i. The first lineage coalesces with the ancestral lineage at time two, just like the second individual lineage.

The trunk of the tree represents the evolution of a genetically unstructured ancestral population. Sequences at the leaves of the tree differ from ancestral sequences at internal nodes through singleton mutations only. Thus ancient variation reflects in the present-day genetic data directly. Ordering samples according to geographic distance from a reference population shows that the UM can be equivalently represented as a comb tree (Figure 1B), similar to the skeleton tree of coalescent-based serial founder effect models [9, 6, 7]. In the UM, each new population is a subset of the source population, and ultimately contains a single lineage for large *n*. The UM does not assume particular migration patterns, and the samples may be randomly scattered in geographic space at arbitrary distance from the geographic origin of their ancestors. A command for the ms program simulating coalescent genealogies under the UM is provided in Appendix A [24].

#### Allele frequency spectrum

Before describing population genetic structure from observed genotypes, we provide some elementary properties of UM genealogies. For a sample of size *n*, the allele frequency spectrum, *ξ*, defined as the frequency of mutations in the sample, can be computed as follows [45]. The probability of a singleton, defined as a mutation being represented uniquely in the sample, is equal the ratio of external branch length to the total length of the genealogy

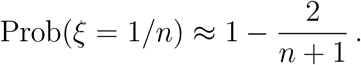

Conditional on non-unique mutations, the frequency of mutation has a uniform distribution

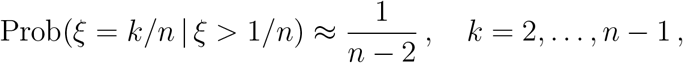

and the unconditional distribution can be described as follows

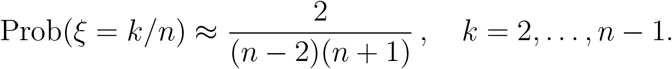

Conditioning on non-unique mutations is equivalent to sampling ancient variation at ancestor nodes in the trunk of the skeleton tree, and the uniform distribution corresponds to the allele frequency spectrum for the ancient samples.

#### Diploid individuals

We limit our discussion to samples of haploid individuals. Diploid individuals could be included in the model by doubling the sample size, considering two individuals from each of the *n* populations. For large *n*, the two lineages in a diploid individual coalesce much before the junction with the trunk of the skeleton tree. For population *k,* the probability of observing an heterozygote locus can be approximated as 4/*n*(*n* + 4), and most genotypes will be homozygote, 0 or 2. The probability of heterogyzosity does not depend on k, reflecting that the UM does not predict any decline of heterogyzosity with respect to a particular origin. In contrast the number of singletons (for haploids) and doubletons (for diploids) can be predicted to increase linearly with the age of populations or with the divergence time from the ancestral lineage.

### 1.2 Eigenvalues and principal components

#### Empirical covariance and Gram matrices

For a data matrix, **Y**, having *n* rows and *L* columns, the first step of a PCA is to transform **Y** so that the mean value of each column is zero, leading to a centered matrix **Z**. The empirical covariance matrix is an *L × L* matrix defined by

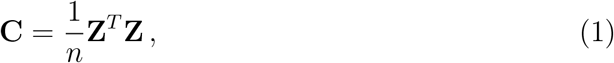

sometimes divided by a factor *n* — 1 for an unbiased estimator of the covariance matrix. Following McVean *[34]*, we consider the *n* × *n* matrix defined by

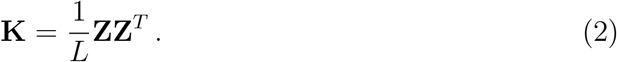

PCA can be performed by computing the eigenvalues and eigenvectors of the matrix **K** as follows

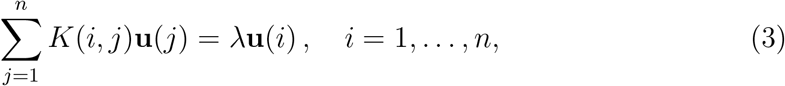

where the *K*(*i,j*)’s are the coefficients of the symmetric matrix **K**. To find approximated solutions, functional forms of eigenvectors will be considered

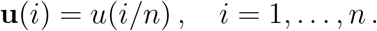

The function *u*(*x*) will be defined on the interval (0,1) as an eigenfunction of an integral operator in an Hilbert space, *L*^2^(0,1). The integral operator will be obtained by considering a continuous approximation of the system of discrete linear equations (3). In the large *n* limit, the continuous approximation will assume that an ancestral lineage could split continuously across time, leading to small daughter populations that evolve independently of the ancestral population. The approximation will be obtained after scaling both sides of equation (3) by a factor *n* or *n*^2^ depending on whether singleton variants are removed or not. For the models considered in our study, the solutions of the finite system of equations (3) can be approximated by the continuous solutions of an integral equation

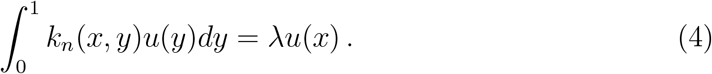

In this equation, *k_n_*(*x,y*) is a continuous kernel that needs to be determined. This will be achieved by considering a coalescent approximation for the coefficients of the Gram matrix as described below.

#### Coalescent approximation

Assuming a neutral coalescent model for the genealogy of a sample of size *n*, a theoretical approximation for the mathematical expectation of the matrix **K** was obtained by McVean [34] as follows

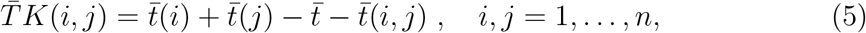

where 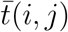 is the average coalescence time of lineage *i* and lineage 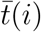 is the average coalescence time of a random lineage with 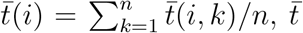 is the average coalescence time of two random lineages, 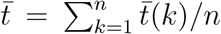, and 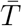 is the average length of the coalescent tree. In those estimates, a *random lineage* is a lineage picked at random in 1, …, *n* from the uniform distribution. Equation (5) is a valid approximation of the sample covariance matrix when the number of genomic loci included in the data matrix is large. For the UM, the average length of the coalescent tree is approximately equal to the length of the skeleton tree. This length is equal to 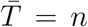 when the singletons are removed from the data matrix, and to 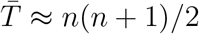 otherwise.

## 3 Main results

The main results of our study are mathematical descriptions of the PCA eigenvalues and eigenvectors for *n* multilocus genotypes sampled from the umbrella model. The first components exhibit wave-like patterns as observed in Cavalli-Sforza et al.’s paper [35, 16, 8]. While Novembre and Stephens showed that sinusoidal patterns arise in principal components when PCA is applied to spatial data [36], our results support the evidence that those patterns may not always be a consequence of spatially structured data. Here, waves result from genetic drift alone, and the hierarchy of population splits from a genetically homogeneous ancestral population without any geographical structure in the sample.

### 2.1 Spectral analysis with singletons removed

This section describes the eigenvalues and eigenvectors of a PCA for a sample of *n* multilocus haploid genotypes after the data matrix has been filtered out for singletons (variants that are uniquely represented in the sample). Removing those rare variants is commonly done in empirical analyses of population genetic structure, often by considering a threshold for the minor allele frequency. Here we condition on non-unique polymorphisms for two reasons. The first reason is aesthetic: the eigenvalues and the principal components have simple exact expressions in the large *n* limit, and they provide direct mathematical evidence for sinusoidal patterns in the PCA. The second reason is a connection to ancient DNA. Removing singletons from present-day samples in the UM amounts to studying samples from their ancestors in the internal branch of the tree. Under the UM, filtering out singletons provides as much genetic information on ancestors as ancient DNA would do, without the shortcomings of the technology.

For non-singleton variants, the average length of the conditioned genealogy is equal to *n*. The covariance matrix is determined by the coalescence times within the ancestral population. More specifically, the average coalescence time for ancestor *i* and ancestor *j* can be described as

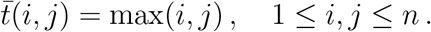

The average coalescence time of ancestor *i* with a random ancestor is equal to

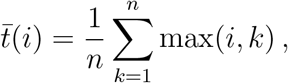

and the average coalescence time of two random ancestral lineages is equal to

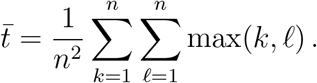

We are now positioned to state our main result, holding for the PCA of a filtered data set. *Let n be a large integer and k* ∈{1,…,*n* — 1}. *For n samples from the UM, the kth largest eigenvalue of the empirical covariance matrix can be approximated as*

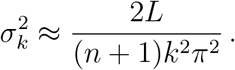

*The kth principal component can be approximated as*

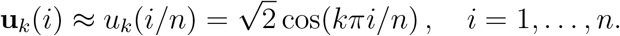

The proof of this result is detailed in Appendix B. It follows from differential calculus arguments based on the approximation of the discrete spectral problem by integral equations. For *x* ∈ (0,1), the continuous eigenvalue problem corresponding to the Gram matrix in equation (2) is

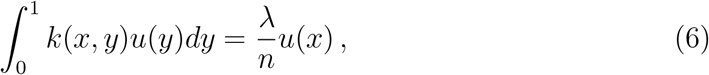

where λ and *u*(*x*) are the unknowns and

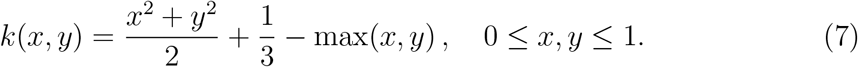

Finding λ and *u*(*x*) is equivalent to solving differential equations of the following type

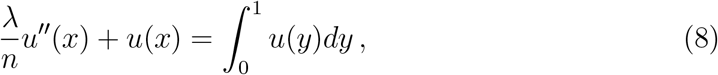

with boundary conditions *u*’(0) = 0, *u*’(1) = 0, and

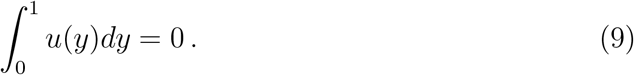

The solutions are provided by the cosine functions and by the values of λ given above (Appendix B). The scaling value in the statement of result, 2*L*/(*n* + 1), is introduced for describing the eigenvalues of the empirical covariance matrix instead of eigenvalues of the Gram matrix. This value represents the number of non-singleton variants in the data matrix. More generally, this value could be replaced by the number of loci in the filtered data matrix. Similar results hold when PCA is extended to data matrices filtered out for doubletons, tripletons or any finite number of classes in the allele frequency spectrum. Minor allele frequency filters, such as considering variants of frequency greater than 5%, could also be analyzed with integral operators. This would require modifying the bounds of the integral, and would lead to less elegant solutions than in the case of singletons.

### 2.2 Connection to ancient DNA and Brownian models

For the data with singleton removed, the PCA of samples obtained from the coalescent genealogies of the UM is similar to the PCA of samples from a random mating population obtained at different time-points in the past (ancient DNA). Here, we describe this result more rigorously, and also establish an equivalence with the Brownian model of drift in allele frequencies *[4]*. Consider a sample of *n* DNA sequences obtained at regularly spaced time points, *t*_1_ = 0, *t*_2_, …, *t*_*n*-1_, *t_n_* = *T*, from a random mating population (*t*_1_ = 0 corresponds to a present-day sample, and *T* is the most ancient sample date). The average coalescence time of two sequences sampled at times *s* and *t* is equal to *τ*_2_(*s, t*) = max(*s, t*) + 1. Thus, average coalescence times are equal to

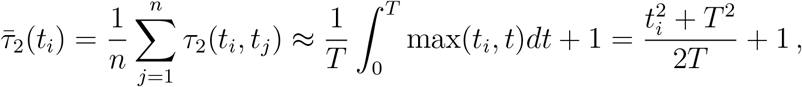

and

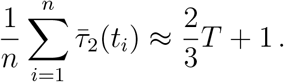

Rescaling time by *T* (*t/T* becomes *t*), the coefficients of the Gram matrix for samples obtained at time *s* and *t* are proportional to

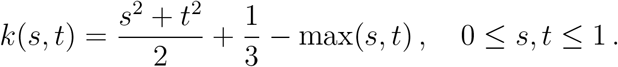

The above kernel corresponds to the kernel arising in the Gram matrix of the data without singletons under the UM. Thus, the principal components of the filtered data are not distinguishable from principal components of data obtained from ancestors sampled at regularly spaced time points in the past. To provide additional interpretation of the Gram matrix coefficients, let us argue that the same kernel arises as a direct consequence of the diffusion approximation in neutral population genetics [27, 28]. More specifically, Cavalli-Sforza and Edwards modelled drift as a Brownian motion, (*B_t_*), 0 ≤ *t* ≤ *T* [4]. For the Brownian motion, the covariance of allele frequencies at time s and t is equal to min(*s, t*). In that case, the Gram matrix can be obtained from the covariance function of the Gaussian process defined as

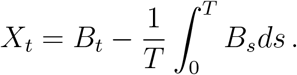

Here the integral term corresponds to the centering of a column in the data matrix. Elementary probabilistic calculus shows that the covariance function of the Gaussian process (*X_t_*) is given by *k*(*s/T,t/T*). Thus, the principal components of the UM are not distinguishable from principal components of Brownian allele frequencies sampled at regularly spaced time points. The result is a justification for the use of a factor model to correct sinusoidal shapes in PCA of ancient DNA [17]. Note that the results also predict that wave-like patterns arise in PC plots obtained from multiple phenotypic values when phenotypes evolve by neutral mutation [32, 14], and provide a way to correct for the distortions caused by temporal sampling.

### 2.3 Spectral analysis with singletons included

This section describes the eigenvalues and principal components for a sample of *n* multilocus haploid genotypes under the umbrella model. All variants are now represented in the sample. In the large *n* limit, the eigenvalues and the principal components have more complex expressions than with filtered data, but their expression still provide mathematical evidence for wave-like patterns in PCs. For present-day samples, the average coalescence time of lineage *i* and lineage *j* can be described as

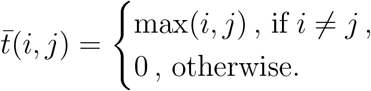

The difference with the case of filtered data is that instantaneous self-coalescence leads to 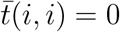 for every individual *i*, a change that complicates the mathematical analysis of the eigenvalue problem for the UM. The average coalescence time of lineage *i* and a random lineage can be described as

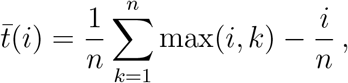

and the average coalescence time of two random lineages is equal to

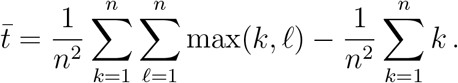

Let *J_α_*(*x*) and *Y_α_*(*x*) denote the Bessel functions of the first and second kind for *α* =1,2 (and for *α* = 0 in a later statement). We formulate our main result for the unfiltered data set below.

*Let n be a large integer value, k ∈ {1,…, n — 1}, and λ_k_ be the kth eigenvalue of the Gram matrix for n samples from the umbrella model. Assume that λ_k_ > 2/n, then λ_k_ can be approximated by the kth largest zero of the function*

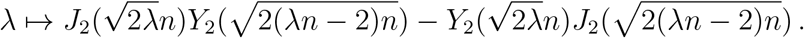

*The kth eigenvalue of the empirical covariance matrix can then be approximated as Lλ_k_/n. The kth principal component,* **u**_*k*_, *can be approximated as*

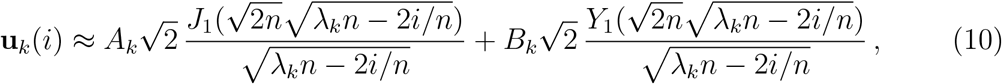

*where B_k_ is defined as*

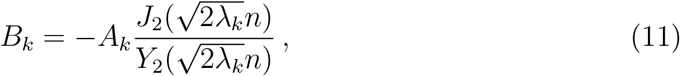

*and A_k_ is a normalizing constant for a unitary eigenfunction (see Appendix C).*

For *n* = 30, the approximation holds for the first three eigenvalues and eigenvectors according to the condition λ_*k*_ > 2/*n*. For *n* = 100, it describes five eigenvalues and eigenvectors, and this number grows to eight eigenvalues and eigenvectors for *n* = 200. For large n, the condition can be verified for about 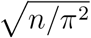 eigenvalues and eigenvectors. As for the case of filtered data, the results were obtained from classical differential calculus arguments. Those arguments are based on the approximation of the discrete spectral problem by an integral equation. For *x* ∈ (0,1), the integral equation is now defined as follows. For *x* ∈ (0,1), we have

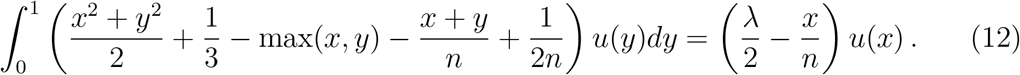

The difference with the analysis of the filtered data set is with the right hand side, which now includes the term equal to –*xu*(*x*)/*n*. The solutions arise from a differential equation of the following type

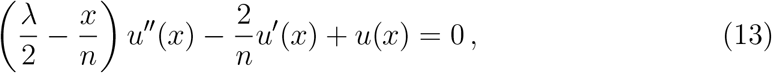

which is similar to equation (8) for large sample sizes, and has the same boundary conditions.

The proof of the result is detailed in Supplementary Text. Although less obvious than for the filtered data, the approximate eigenvectors still provides evidence of wave-like patterns arising in PC plots. As the simulation study will show, those eigenvectors capture wave patterns more complex than those obtained with the cosine functions.

## 3 Simulations and data analysis

In order to check the accuracy of continuous approximations of PCA eigenvalues and eigenvectors, we performed coalescent simulations of haploid genotypes under the umbrella model. In this section, the results were reported for *n* = 100 populations and samples. Results for *n* = 30 populations (and samples) could be found in the supplementary materials. The number of populations (*n* = 30, 100) was chosen to be of the same order of magnitude as in our analysis of human genotypes from the Simons Genome Diversity Project [33]. Simulations of the UM were performed with the program `ms`. PCA was performed in the `R` programming language with the command `prcomp`. First, we compared the eigenvalues and eigenvectors of a PCA for genotypes filtered out for singletons. We recall that an interesting property of the model is that removing singletons amounts to observing ancestral variants, corresponding to the evolution of a single ancestral population sampled at regular time points. Then we compared the eigenvalues and eigenvectors of the PCA for the unfiltered data set, in which the number of singletons in a particular individual sample reflects the age of its ancestor in the internal branch of the skeleton tree.

### Singletons removed

PCA eigenvalues, corresponding to the variance of projected samples along each PC 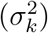, were compared to their approximation obtained from equation (8). According to our theoretical result, 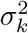 should be approximately equal to *L*’/(*π*^2^*k*^2^), where *L*’ is the number of loci in the genotype matrix (*L*’ ≈ 2*L*/(*n* + 1)). We found that the approximations of PC eigenvalues were highly accurate, and they were also close to numerical eigenvalues obtained by solving the discrete system of linear equations (2) based on McVean’s matrix (Figure 2, top panels). The results provided evidence that the empirical matrix was well-approximated by its expected value – a result that holds by the law of large numbers –, and that the integral approximation was also accurate. Next we checked that PCs exhibited wavelike patterns as predicted by the infinite-size theory (Figure 2). For the four first PCs, we found that the simulated PCs were hardly divergent from their continuous approximations by cosine functions, 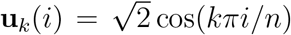, for *i* = 1,…,*n*. The PC plots, showing PC_k+1_ as a function of PC_k_, also displayed wave-like patterns (Figure 3). The PC1-PC2 plot exhibited a symmetric arch, whereas the PC2-PC3 plot exhibited a fish-like pattern. Higher PC plots exhibited patterns similar to Lissajous curves. The results for *n* = 30 were similar to those obtained with *n* = 100, and showed that the number of populations does not need to be very large for the asymptotic theory to apply (Figure S1 and Figure S2). The PC plots also looked like those obtained in simulations of one-dimensional stepping-stone models [36]. The similarity was not expected a priori, because the covariance matrix of the steppingstone model, which has a stationary block-Toeplitz structure, differs from the non-stationary structure of the umbrella model. Whatever the value of n, the remarkable fact is that the sinusoidal patterns arise without any geographic structuring, but because a genetically homogeneous founder population gives birth to many small drifting subpopulations through its temporal evolution.

**Figure 2.**
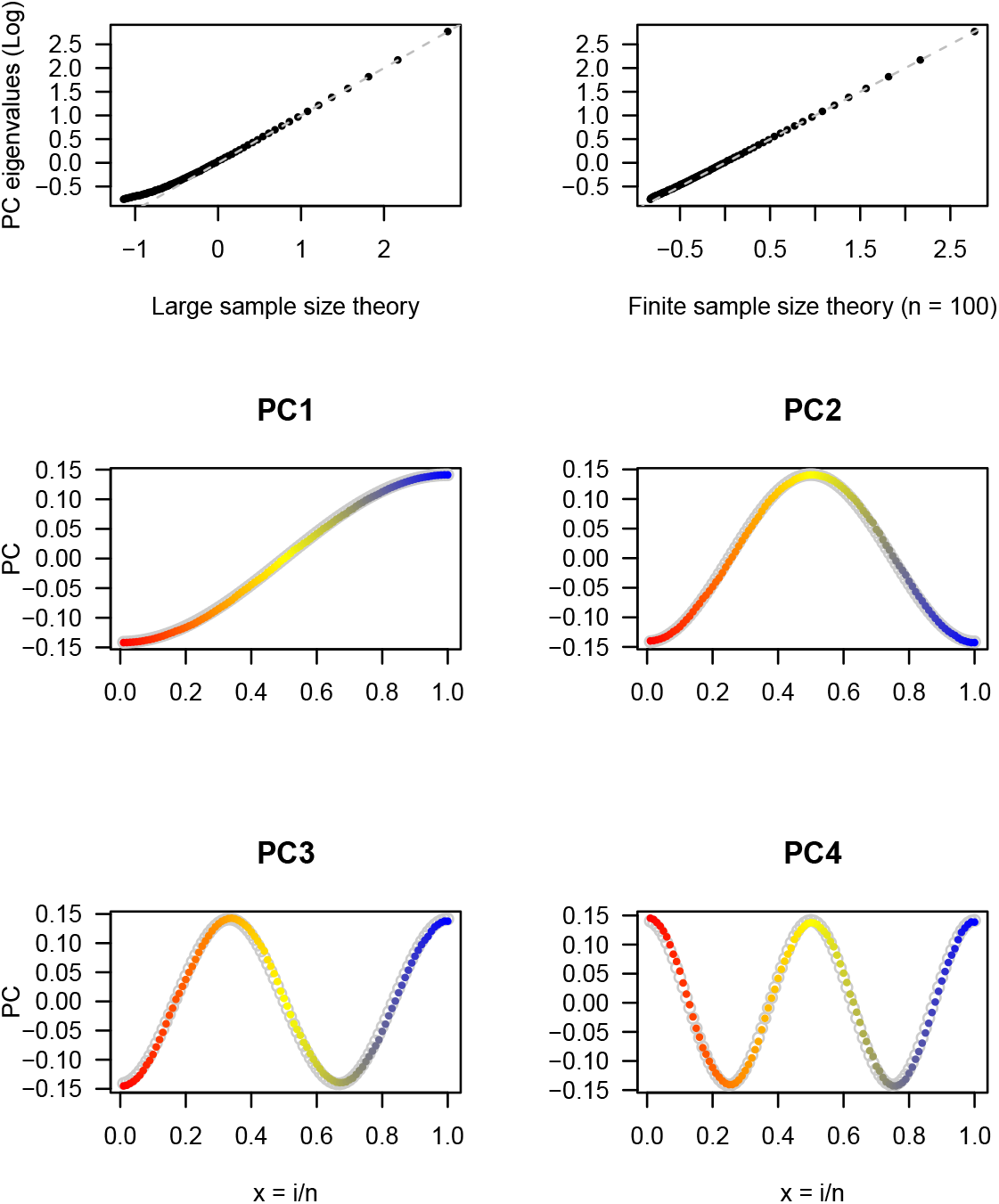
Approximation of PC eigenvalues and eigenvectors for filtered data from the UM. Simulation of the umbrella model performed for *n* = 100 haploid individuals and *L’* = 5,938 single nucleotide polymorphism loci obtained after removing singletons. *Top panels:* Approximations of PC eigenvalues by *L’/π*^2^*k*^2^ (left) and by the numerical values obtained from the discrete system of linear equations (right). Eigenvalues are displayed on a base 10 logarithm scale. *Bottom panels:* Approximations of PC eigenvectors by their continuous approximations, 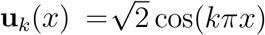, for *x* = *i/n, k* = 1 to 4. Eigenvectors are represented in red (most recent samples) to blue (most ancient samples) colors. Continuous approximations are represented by grey circles.

**Figure 3.**
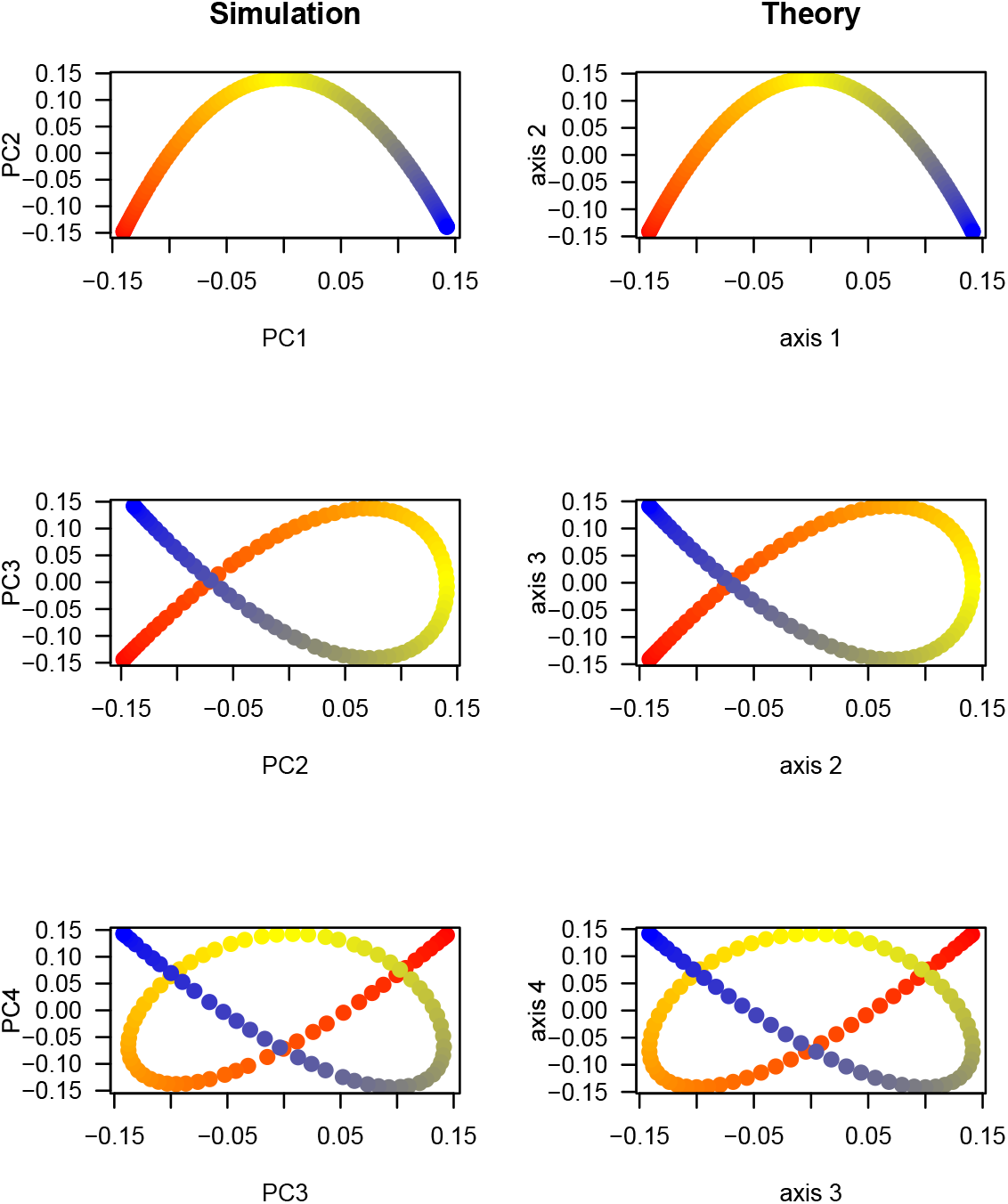
Wave patterns in PC plots for filtered data. Simulation of the umbrella model performed for *n* = 100 haploid individuals and *L*’ = 5, 938 single nucleotide polymorphism loci obtained after removing singletons. *Left column*: PC plots for the simulated data. *Right column*: Continuous plots derived from the cosine approximations for *x* = *i/n* and *k* = 1 to 4. Eigenvectors are represented in red (most recent samples) to blue (most ancient samples) colors.

### Singletons included

For unfiltered data from the UM, the contribution of singletons to genetic variation modified the shapes of PCs, and thereby, the patterns of population structure changed significantly (Figure 4, Figures S3–S4). For the first five PCs, we found that the approximations of PC eigenvalues were highly accurate, and again very close to numerical eigenvalues obtained from solving the discrete system of linear equations (Figure 4 top left panels, and Figure S5 for *n* = 30). The PCs exhibited wave-like patterns as predicted by the Bessel functions obtained from theoretical approximation by the integral operator (Figure 4, Figure S3). The amplitude of oscillation of these functions varied with the age of the individual ancestor, corresponding to the PC ranking. The PC plots exhibited shapes that were not observed in simulations of one-dimensional stepping-stone models. The PC1-PC2 plot still exhibited an arch pattern, but the arch had an asymmetrical shape, with a more pronounced asymmetry for smaller *n* (Figure S3 and Figure S5). The first PCs remained close to those obtained for the filtered data set, showing that the first axes of variation are mainly influenced by genetic drift in the ancestral population. However, the subsequent PCs exhibited patterns that were unseen in the PCA of the filtered data set. PCs associated with smaller variances were flat until the age of the ancestor, exhibited a large amplitude oscillation, and then returned back to zero quickly (Figure S4). Those shapes made PC plots difficult to display graphically, and only a few first PCs could be usefully interpreted, especially in the smaller *n* case.

**Figure 4.**
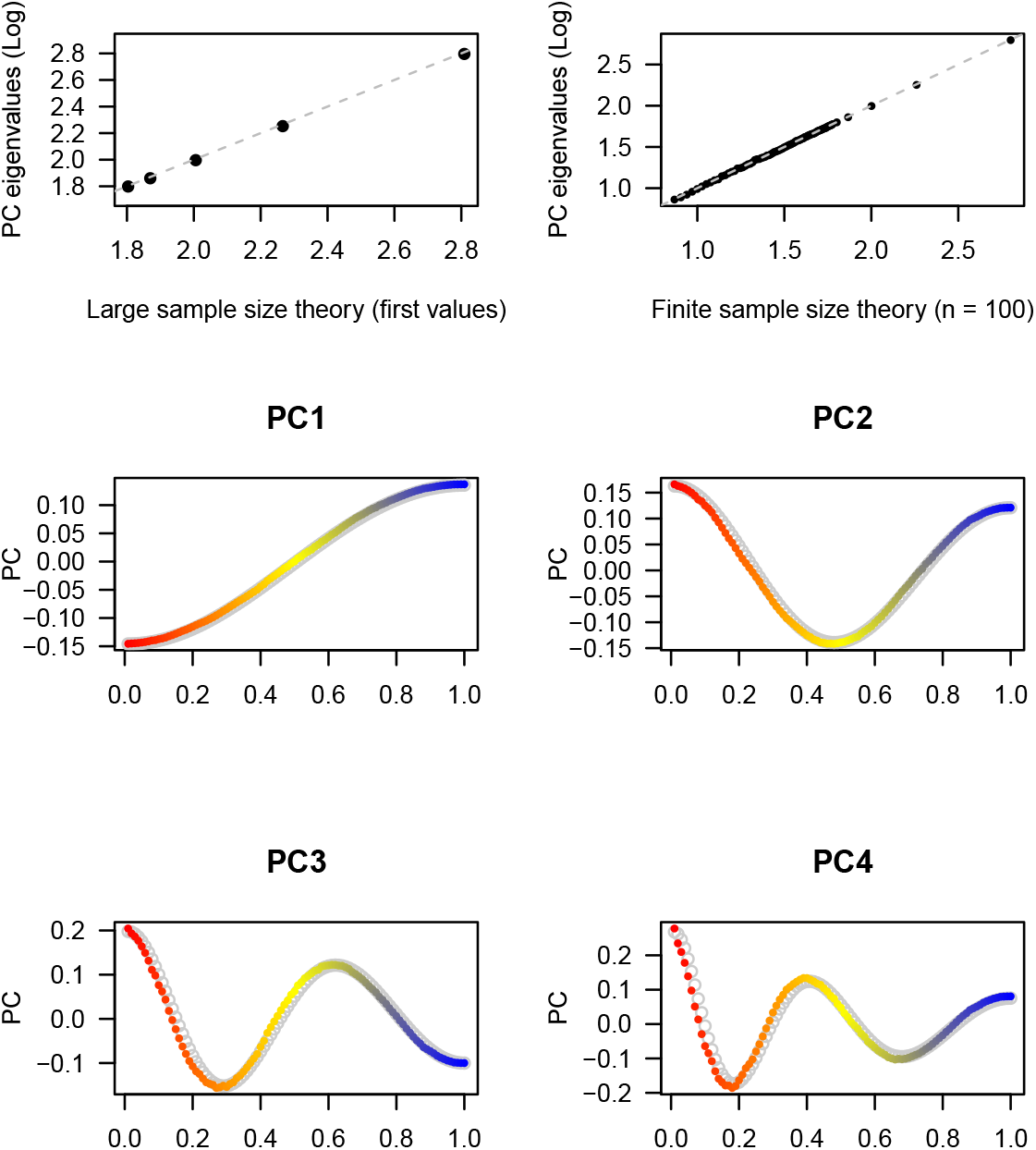
Approximation of PC eigenvalues and eigenvectors for unfiltered data. Simulation of the umbrella model performed for *n* = 100 haploid individuals and *L* = 300, 000 single nucleotide polymorphism loci. *Top panels*: Approximations of the five first PC eigenvalues by *L*λ_*k*_/*n*, where *λ_k_* is obtained from the continuous approximation (left), and by the numerical values obtained from the discrete eigenvalue problem using McVean’s equations (right). Eigenvalues are displayed on a base 10 logarithm scale. *Bottom panels:* Approximations of PC eigenvectors by their continuous Bessel approximations, **u**_*k*_(*x*), for *x* = *i/n, k* = 1 to 4. Eigenvectors are represented in red (most recent samples) to blue (most ancient samples) colors. Continuous approximations are represented by grey circles.

### Human data analysis

To evaluate the relevance of the umbrella model to biological data, we analyzed human genomic data for fifty-nine individuals originating from *n* =30 human populations in Asia and in the Americas [33]. The individual identifiers and the geographic coordinates of the sampled populations were given in Table S1, and displayed in Figure 5. The study of those populations was motivated by the settlement of the Americas, for which the prevalent models outline Asian migration from the Beringia land bridge and subsequent dispersal of the ancestral population throughout the Americas [21, 44]. A subset of around 555k SNPs was drawn from the whole genome sequence data, after filtering out missing data and variants with minor allele frequency less than 5%. To keep the dimension of the data similar to those used in the simulation study, we checked that the results obtained with 10k SNPs were robust to subsampling and similar to those presented in Figure 5 (Figure S6).

**Figure 5.**
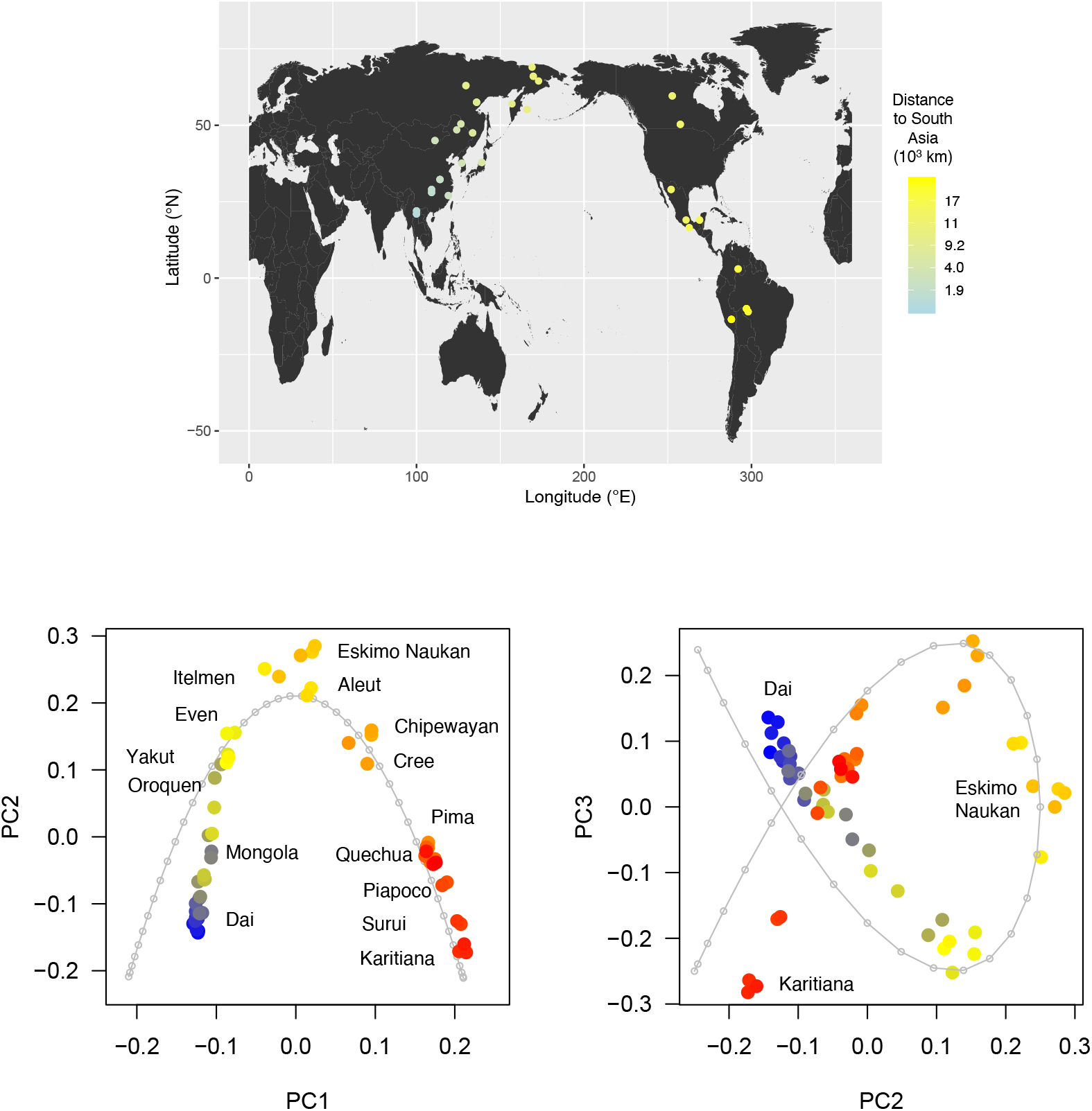
Human data analysis. *Top panel:* Geographic coordinates of 59 samples from the Simons Genome Diversity Project (Table S1). The color gradient reflects the geographic distance to China. *Bottom panel:* PC plots for the genotype data (554,716 SNPs). The color gradient reflects coalescent time. Blue: Dai (21° N, 100° E), Red: Karitiana (Brazil, —13.5° N, 348° E). The grey lines represent the predictions from the umbrella model with *n* = 41 populations (PC1-PC2 plot) and *n* = 32 populations (PC2-PC3 plot).

The first two eigenvectors of the PCA displayed a horseshoe pattern for which the most extreme populations were the Dai population of China (21° N, 100° E) and the Karitiana population of Brazil (—13.5° N, 348° E). The observed arch agreed with the predictions of a UM for which the most ancient populations were Asian and the most recent ones were South American (Figure 5). Nevertheless, a constant ancestor population size in the UM did not allow a perfect fit to the data. For example, the number of present-day populations was overestimated by the UM (*n* = 41). In approximations of PC2 and PC3, the number of populations in the UM (*n* = 31) was closer to the number of present-day populations in the samples (*n* = 30), and the second and the third eigenvectors of the PCA displayed a fish-like pattern that agreed with the predictions of the random drift model.

To evaluate whether the sinusoidal patterns observed in PCA also arise in ancestry estimates obtained with other methods, we performed an analysis of population genetic structure with a program that reproduces the results of STRUCTURE [41, 20]. The STRUCTURE analysis identified two ancestral genetic groups corresponding to the Asian and South American populations in the PC1 curve (Figure 6). Asian genetic ancestry measured with STRUCTURE (blue color in Figure 6) exhibited a very high correlation with the first PC *(r* = 99.8%). For the examples of Aleut and Eskimo populations, a direct interpretation of the admixture estimates contradicts recent evidence that migrations connected to the peopling of the Aleutian islands are genetically linked to a single Siberian source related to Paleo-Eskimos [15]. The correspondence between PCA and STRUCTURE has been outlined in several studies [37, 12]. Although STRUCTURE interprets the data in terms of admixture coefficients, drift alone might be the most parsimonious explanation for the observed clinal pattern, and that explanation does not resort to any complex spatial processes.

**Figure 6.**
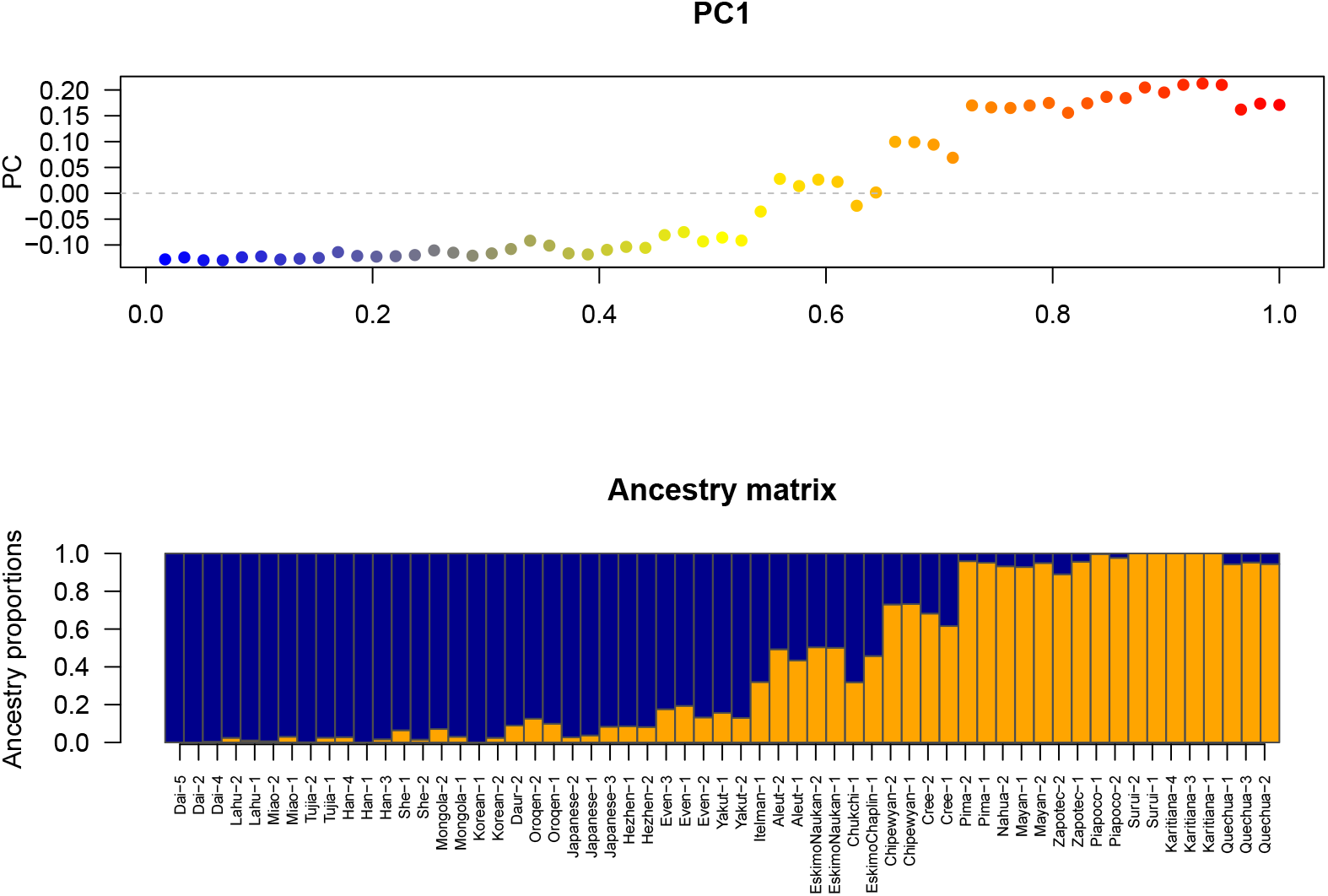
Comparison of PCA and STRUCTURE for human samples. For 59 Asian and American samples from the Simons Genome Diversity Project, PC1 displayed a sinusoidal shape that agreed with the UM (*top panel*). For *K* = 2, the wave pattern was also reflected in the admmixture plot (*bottom panel*, Pearson correlation r = 99.8%. The color gradient reflects the geographic distance to China. Blue: Dai (21° N, 100° E), Red: Karitiana (Brazil, —13.5° N, 348° E).

## 4 Conclusion

This study provided a detailed mathematical analysis of the eigenvalues and eigenvectors obtained from the principal component analysis of a diffusion model of genetic variation. The model was based on hierarchy of population splits from a genetically homogeneous population without any geographical structure in the sample. Our results described continuous approximations of PC eigenvectors for large sample size. When singletons are removed from the genetic data, the PCA eigenvectors took simple mathematical expressions, defined as cosine functions of increasing periodicity. For unfiltered data sets, the first PC eigenvectors could be approximated by Bessel’s functions that exhibit complex wave-like patterns.

Novembre and Stephens showed that sinusoidal patterns arise in principal components generally when PCA is applied to spatial data *[36]*. The waves were interpreted as mathematical artifacts that can be observed when covariance between locations decays with spatial distance. Their study suggested that the artificial patterns could be amplified by spatial interpolation methods used to display PC eigenvectors as geographic maps. Our results support the alternative evidence that wave-like patterns could arise from genetic drift only, and may not be artificial outcomes of spatially structured data.

Several studies have argued that human genetic variation agrees with a model of isolation-by-distance, whereas others see a role for long-range migrations and bottlenecks [25]. Models of recurrent bottlenecks start with a single source population [9, 6], and consider a series of migration similar to the stepping-stone principle. The source population sends a subset of its individuals to migrate outward and found a new population, which grows and sends out migrants to form the next population. The founding process is iterated until a predefined number of populations have been founded. Although the principal components of human genetic variation considered here agreed with a UM, the model is very different from a stepping-stone model or a serial founder effect model. Perhaps the simplest metaphor of the UM in a spatial context would be as a random mating population moving in geographic space, leaving small daughter populations at every visited location. In this process, a new population is not a subset of a previous population in the expansion, but a subset of the source population itself.

Regarding the implication of the results for human data, we showed an example of complex arch and fish-like patterns observed PC plots when studying the migration of an Asian ancestral population to the Bering Straits and its dispersal throughout the Americas. Although the PC eigenvectors reproduced patterns observed in a one-dimensional equilibrium isolation-by-distance model, this multiple-island model represents an unlikely explanation for the genetic data [15]. In addition, analysis of population structure based on STRUCTURE-like algorithms could lead to misleading estimates of admixture coefficients in Beringian populations. The confounding effect of hierarchical population splits on admixture estimates was confirmed by coalescent simulations (Figure S7, see [29] for how to misinterpret STRUCTURE results). Thus STRUCTURE-like ancestry estimation algorithms did not lead to better explanations than the sinusoidal patterns obtained from the PCA of the human data. Our study shows that interpretation of population genetic data in terms of geographic expansions is very difficult, because models with opposite features can provide similar predictions of population structure. Genetic drift provides a simple null-model that one usually attempts to reject by statistical hypothesis testing. Principal components of the genetic data, which summarize information on population genetic structure, population trees, admixture and inbreeding coefficients cannot reject models of very low complexity such as the umbrella model in this analysis.

## Appendices

### Appendix A. Simulation of umbrella model

Simulations of the umbrella model corresponded to the following ms command [24]

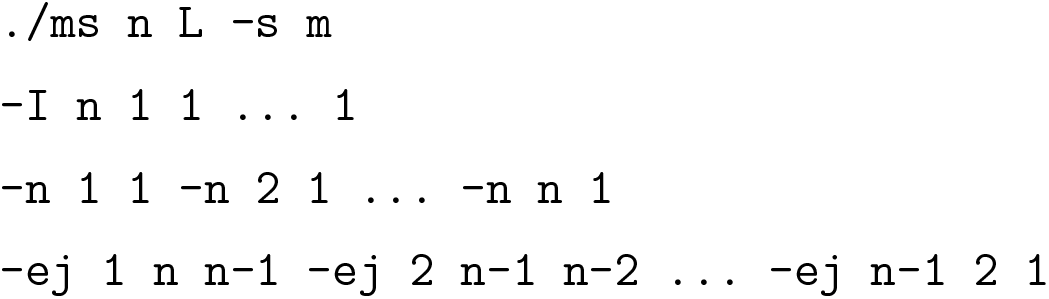

### Appendix B. Proof of the approximation result for filtered data sets

According to [34], a theoretical approximation for the mathematical expectation of the matrix **K** is defined by equation (5). For non-singleton variants, the average total branch length in the tree is equal to *n* and the average coalescence time for ancestor *i* and ancestor *j* is equal to the maximum between *i* and *j*. To find the eigenvectors and eigenvalues of 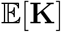, we studied the spectrum of matrix 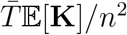, which is equal to 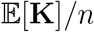. We looked for an eigenvector **u** and his corresponding eigenvalue *μ* such that 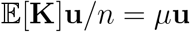. We set λ = *nμ*, so that we have 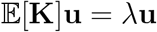. To solve the last equation, we sought solutions of the form **u**(*i*) = *u*(*i/n*), *i* = 1,…, *n*, where *u*(*x*) is a continuous function. With this notation, the spectral equations could be rewritten as

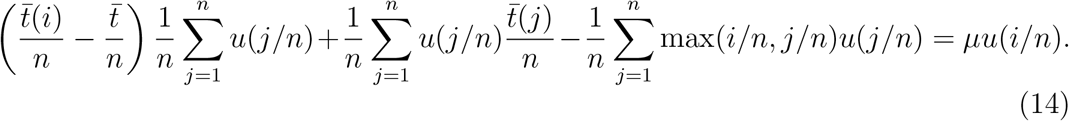

Approximating the Riemann sums by integrals, we obtained

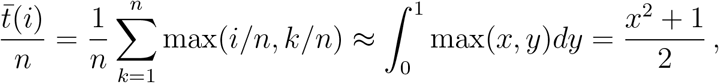

and

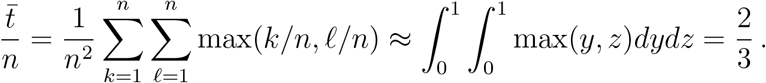

With similar approximations for the other terms in equation (14), the discrete eigenvalue problem becomes a continuous eigenvalue problem

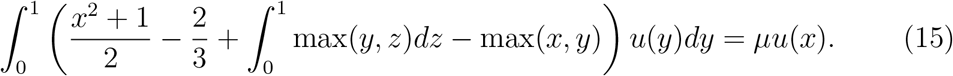

Simplifying (15), we obtained

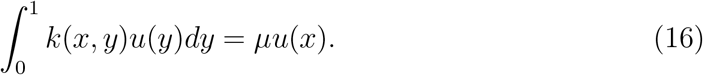

where

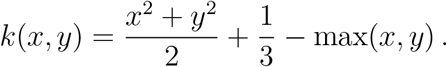

By a direct integration, we found that

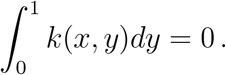

Integrating on both sides of equation (16), led us to

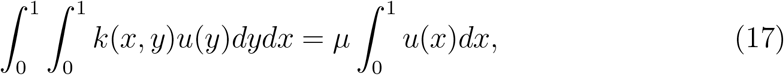

and exchanging order of integrals, we obtained

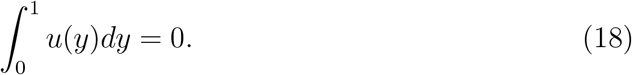

Therefore, equation (15) becomes

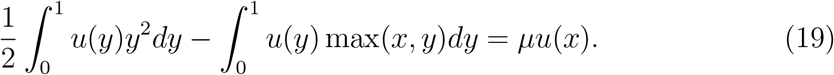

Differentiating (19) yielded

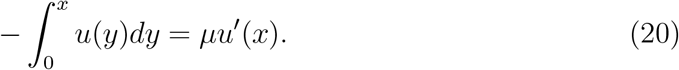

A second differentiation led to the following differential equation

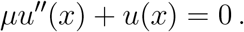

The solutions of the differential equation have the following form

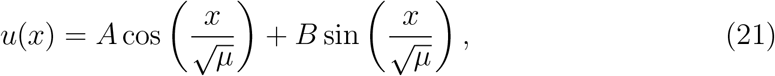

where *A* and *B* are constant values determined by the boundary conditions. Setting *x* = 0 in equation (20) gave *u*’(0) = 0 which led to *B* = 0. Setting *x* =1 gave *u*’(1) = 0, which implies that the eigenvalues were equal to *μ_k_*= 1/(*k*^2^*π*^2^), for *k* = 1, …, *n* – 1. Thus the normalized eigenvectors and their corresponding eigenvalues had the following form

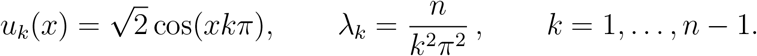

### Appendix C. Normalizing constant for the unfiltered data analysis

The normalizing constant for the unfiltered data analysis is defined by

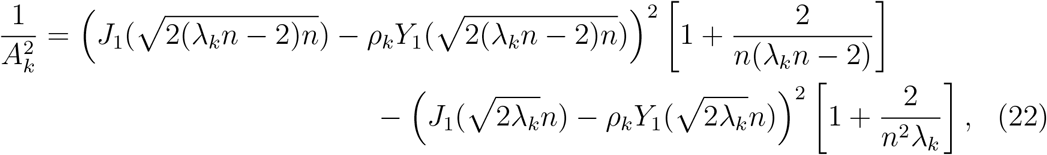

where 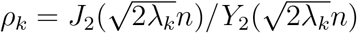.

## Supplementary materials

**Figure S1.**
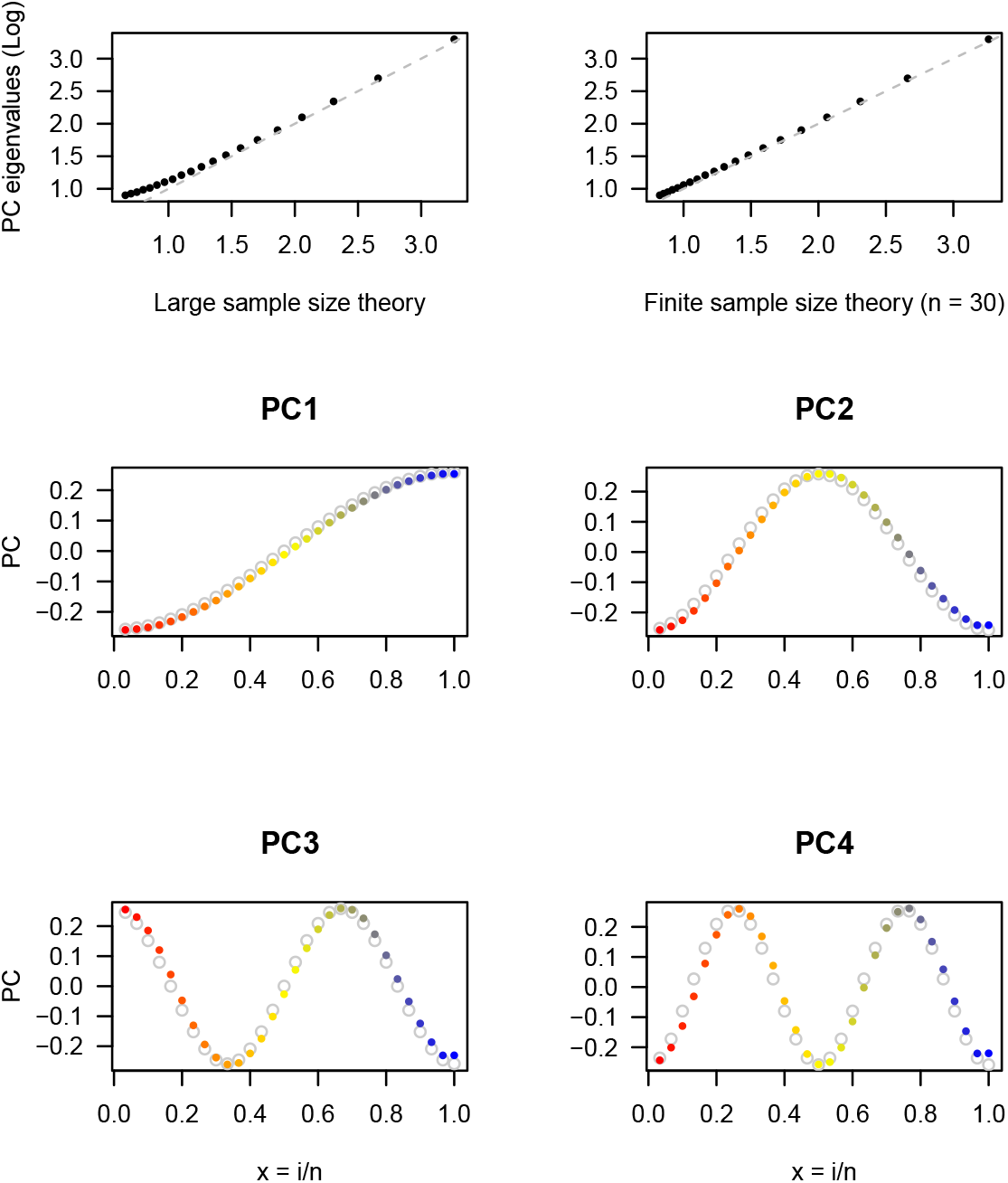
Approximation of PC eigenvalues and eigenvectors for filtered data from the UM (n = 30). Simulation of the umbrella model performed for *n* =30 haploid individuals and *L* = 18,002 filtered single nucleotide polymorphism loci. *Top panels:* Approximations of PC eigenvalues by *Lλ_k_/n* = *L/π*^2^*k*^2^ (left) and by the numerical values obtained from the discrete eigenvalue equations based on McVean’s equation for *n* = 30 (right). Eigenvalues are displayed on a base 10 logarithm scale. *Bottom panels*: Approximations of PC eigenvectors by their continuous approximations, 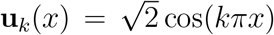, for *x* = *i/n, k* = 1 to 4. PC eigenvectors are colored from red (most recent samples) to blue (most ancient samples) and the continuous approximations are represented by grey circles.

**Figure S2.**
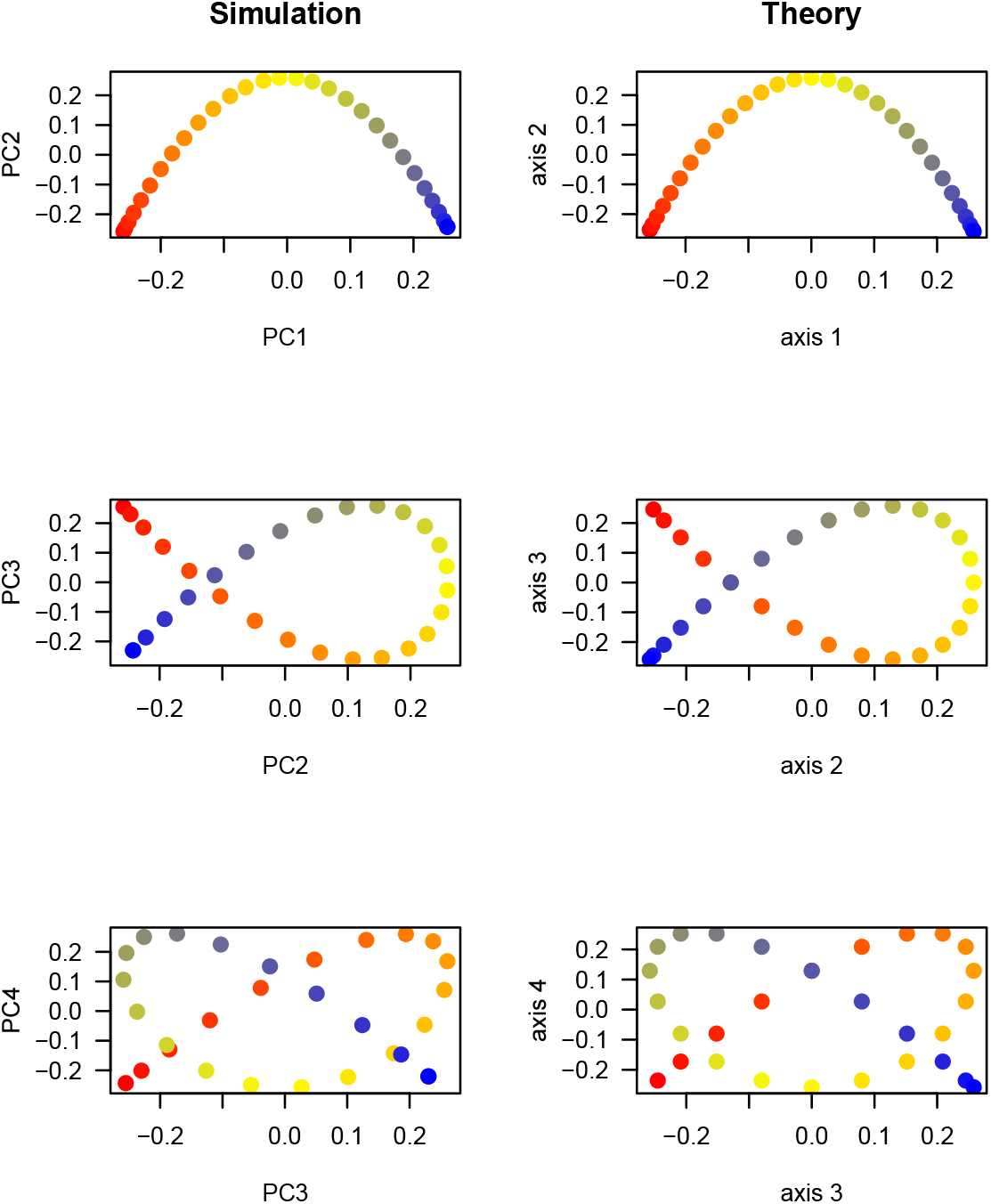
Wave patterns in PC plots for filtered data (n = 30). A simulation of the umbrella model was performed for *n* = 30 haploid individuals, and resulted in *L* = 18, 002 unfiltered single nucleotide polymorphism loci. *Left column:* PC plots for the simulated data. *Right column:* Continuous plots derived from the asymptotic theory for *x* = *i/n* and *k* = 1 to 4. Eigenvectors are represented in red (most recent samples) to blue (most ancient samples) colors.

**Figure S3.**
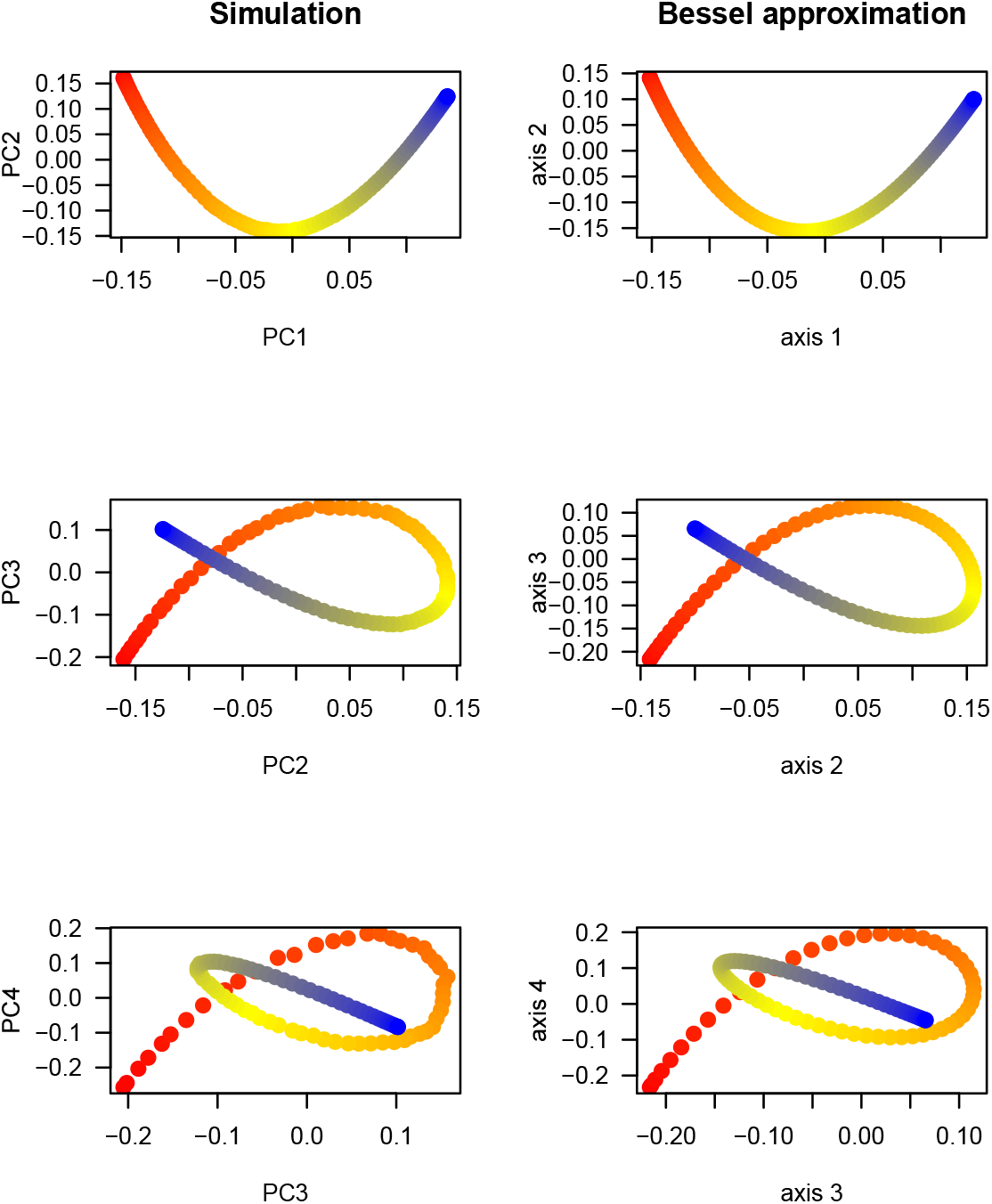
Wave patterns in PC plots for unfiltered data (n = 100). Simulation of the umbrella model performed for *n* = 100 haploid individuals and L = 300, 000 unfiltered single nucleotide polymorphism loci. *Left column:* PC plots for the simulated data. *Right column*: Continuous plots derived from the asymptotic theory (Bessel approximations). Eigenvectors are represented in red (most recent samples) to blue (most ancient samples) colors.

**Figure S4.**
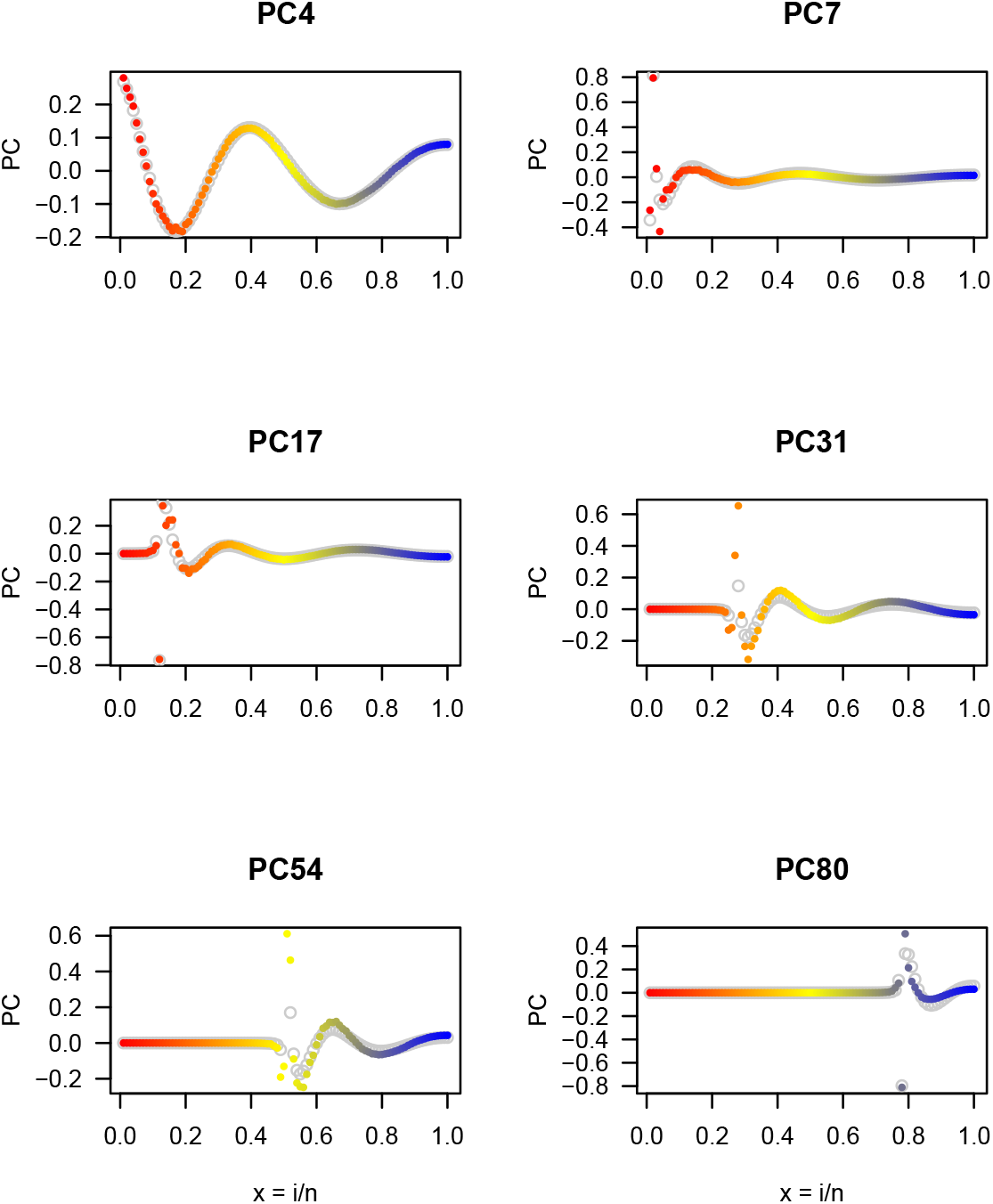
PC eigenvectors for unfiltered data (n = 100). Simulation of the umbrella model performed for *n* = 100 haploid individuals and *L* = 300, 000 unfiltered single nucleotide polymorphism loci. Six PC eigenvectors for the simulated data and their prediction from the discrete system of equations. Eigenvectors are represented in red (most recent samples) to blue (most ancient samples) colors.

**Figure S5.**
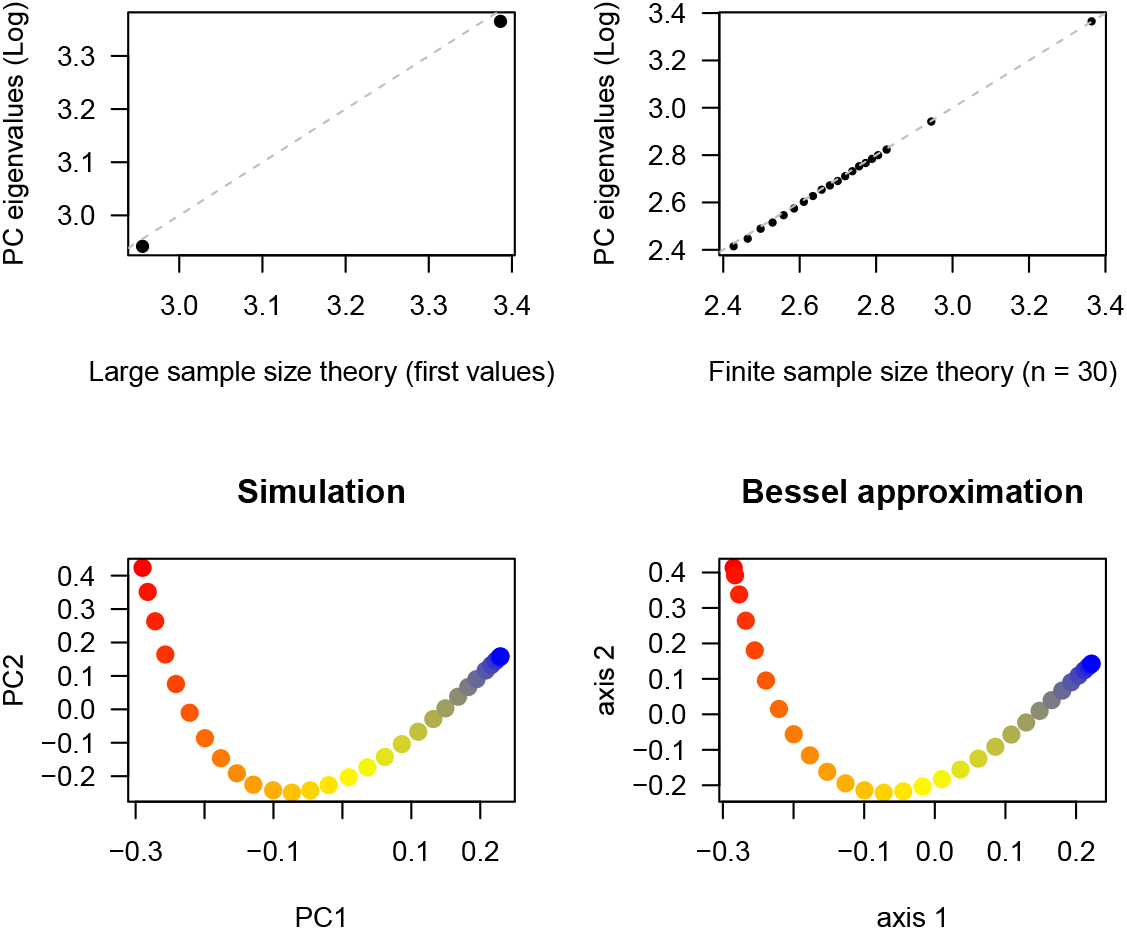
Approximation of PC eigenvalues and eigenvectors for unfiltered data (n = 30). Simulation of the umbrella model performed for *n* = 30 haploid individuals and *L* = 300, 000 unfiltered single nucleotide polymorphism loci. *Top panels:* Approximations of the first PC eigenvalues by *Lλ_k_/n* (continuous approximation) (left) and by the numerical values obtained from the discrete eigenvalue equations for *n* = 30 (right). Eigenvalues are displayed on a base 10 logarithm scale. *Bottom panels*: PC plots for the simulated data and continuous plots derived from the asymptotic theory. Eigenvectors are represented in red (most recent samples) to blue (most ancient samples) colors.

**Figure S6.**
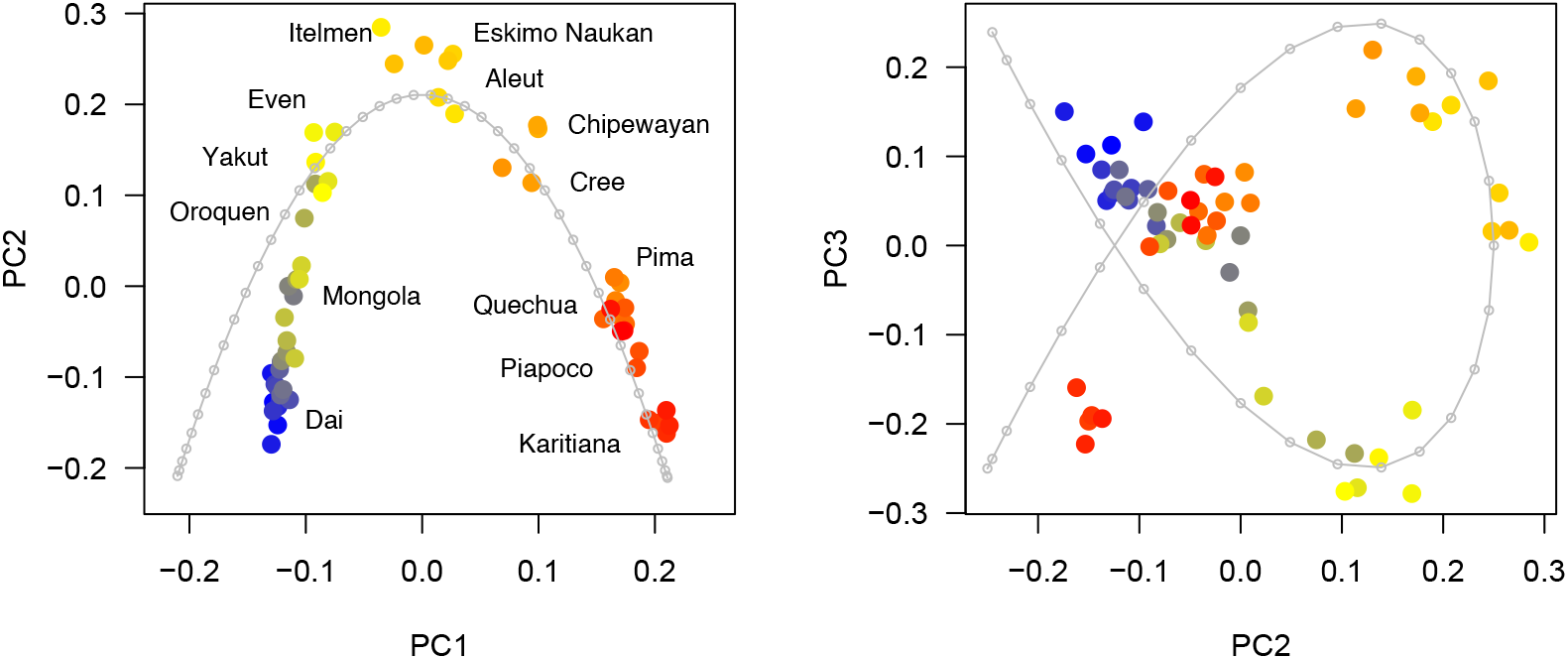
Human data analysis. PC plots for 10k filtered genotypes from 59 individuals from the SGDP. The color gradient reflects the geographic distance to China. Blue: Dai (21° N, 100° E), Red: Karitiana (Brazil, −13.5° N, 348° E). The grey lines represent the predictions from the umbrella model with *n* = 41 populations (PC1-PC2 plot) and *n* = 32 populations (PC2-PC3 plot).

**Figure S7.**
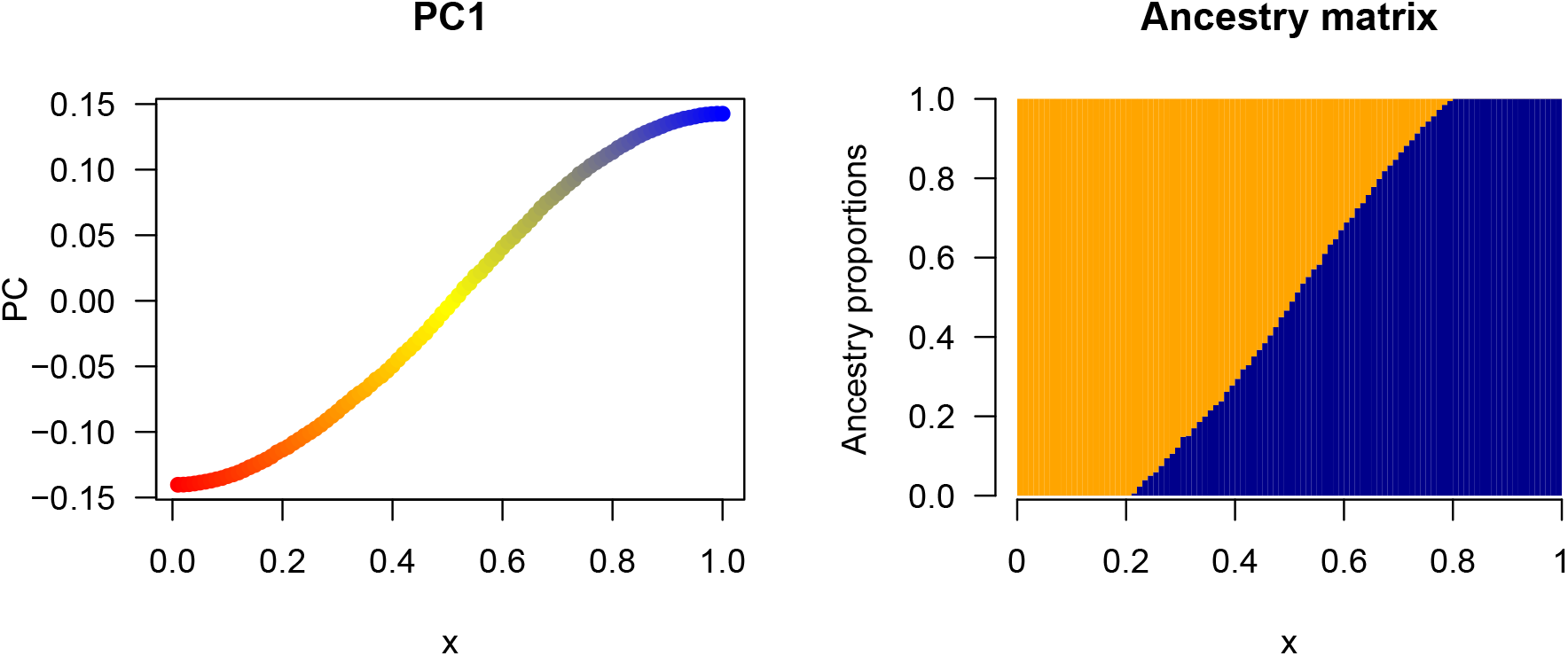
Ancestry coefficients under the UM. Simulation of the umbrella model performed for *n* = 100 haploid individuals and L = 5, 938 filtered single nucleotide polymorphism loci. *Left*: First principal component. *Right*: Ancestry coefficient estimates from a STRUCTURE-like program run for *K* = 2.

**Table S1.**
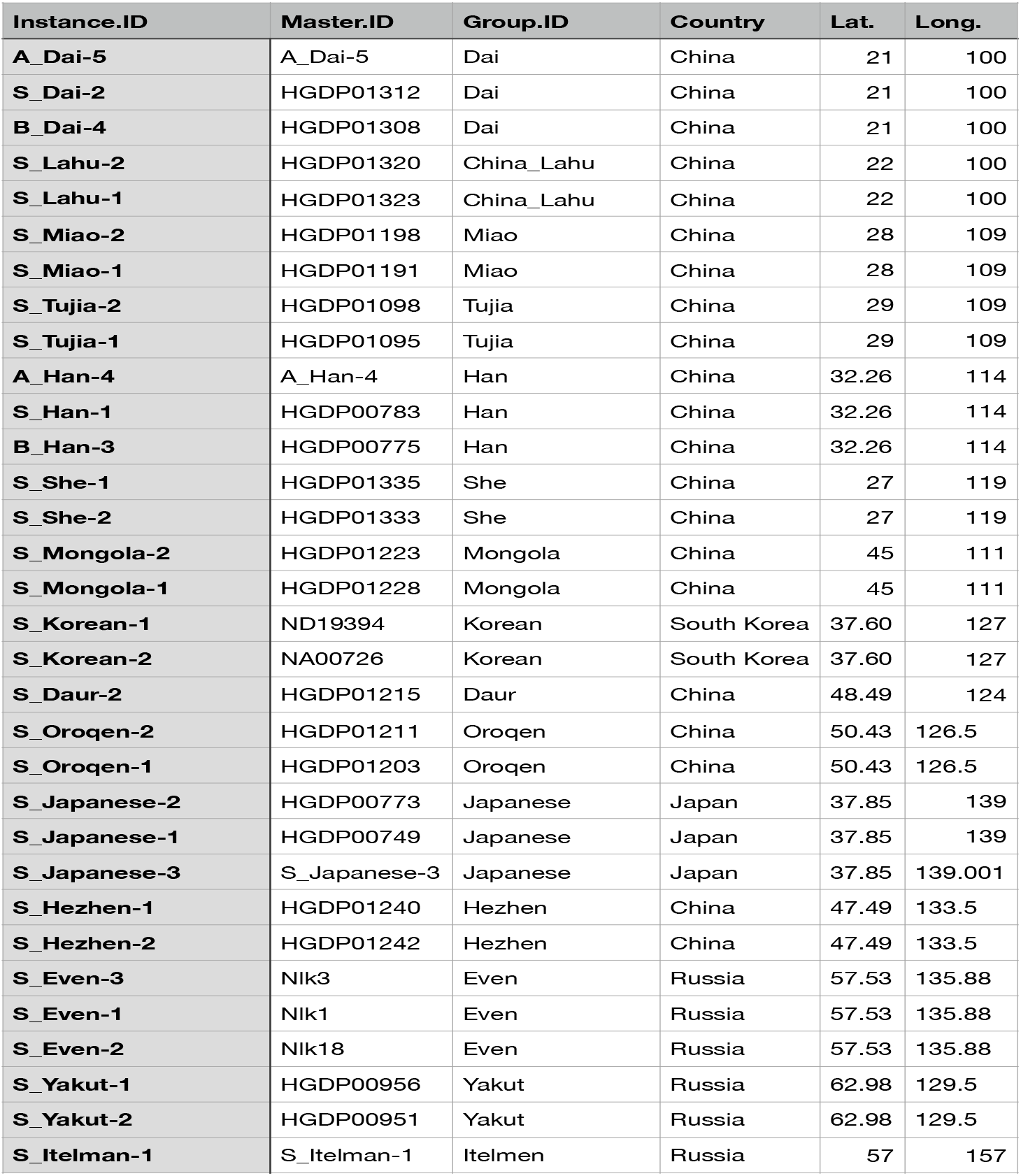
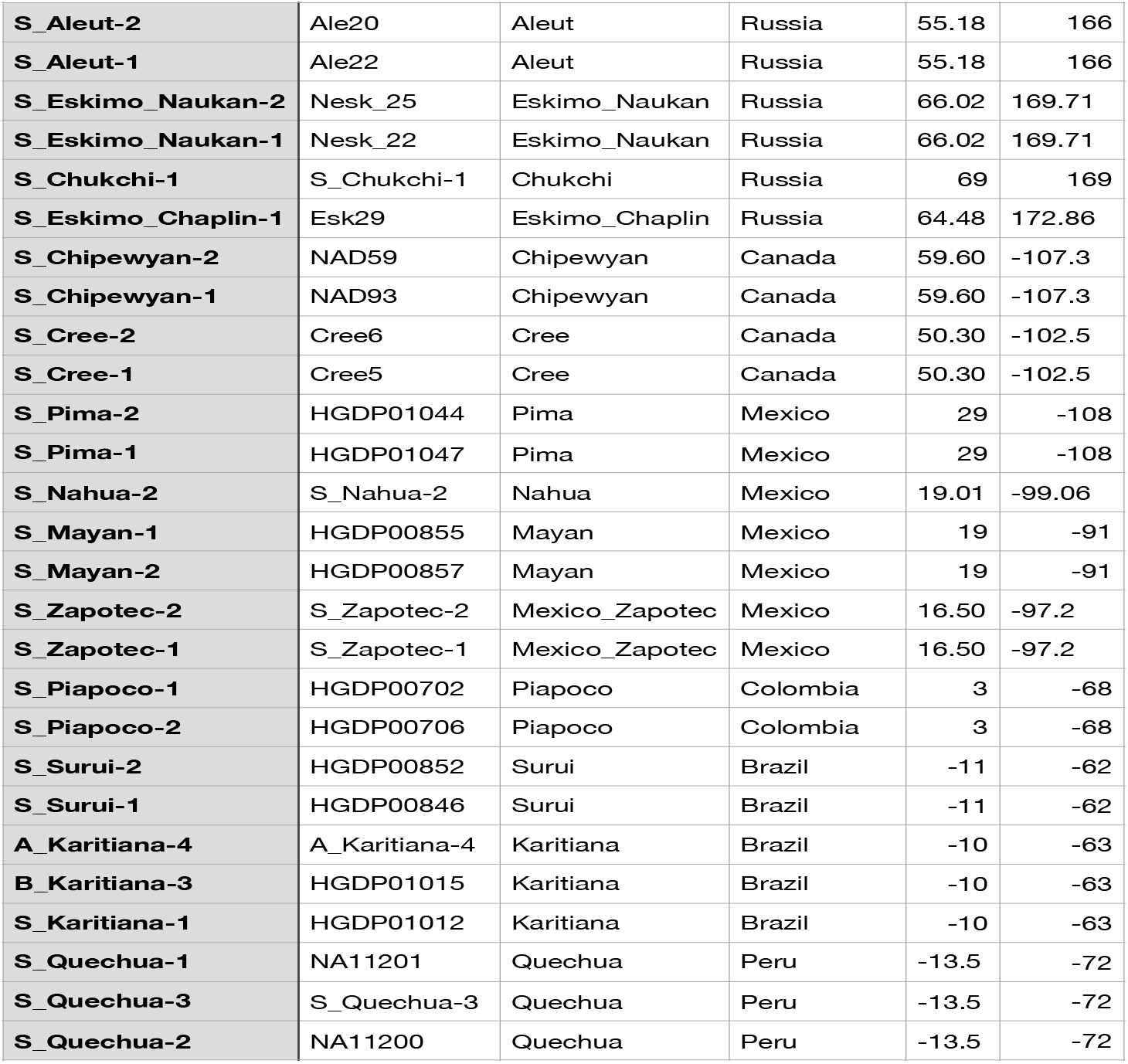
List of human samples used in the study.

