## Supplementary text for "Theoretical Analysis of Principal Components in an Umbrella Model of Intraspecific Evolution"

### 1 Spectral analysis of unfiltered samples from the umbrella model

In this section, we prove the following results.

*Let $n$ be a large integer value, $k \in \{1, \ldots, n\}$, and $\lambda_k$ be the $k$th eigenvalue of the Gram matrix for $n$ samples from the umbrella model. Assume that $\lambda_k > 2/n$, then $\lambda_k$ can be approximated by the $k$th largest zero of the function*

$$\lambda \mapsto J_2(\sqrt{2\lambda}n)Y_2(\sqrt{2(\lambda n - 2)n}) - Y_2(\sqrt{2\lambda}n)J_2(\sqrt{2(\lambda n - 2)n})\,.$$

*The $k$th principal component, $\mathbf{u}_k$, can be approximated as*

$$\mathbf{u}_k(i) \approx A_k\sqrt{2}\,\frac{J_1(\sqrt{2n}\sqrt{\lambda_k n - 2i/n})}{\sqrt{\lambda_k n - 2i/n}} + B_k\sqrt{2}\,\frac{Y_1(\sqrt{2n}\sqrt{\lambda_k n - 2i/n})}{\sqrt{\lambda_k n - 2i/n}}\,, \qquad (1)$$

*where $B_k$ is defined as*

$$B_k = -A_k\frac{J_2(\sqrt{2\lambda_k}n)}{Y_2(\sqrt{2\lambda_k}n)}\,, \qquad (2)$$

*and where $A_k$ is a normalizing constant for a unitary eigenfunction (see Appendix C).*

A theoretical approximation for the mathematical expectation of the Gram matrix $\mathbb{E}[\mathbf{K}]$ was obtained in Ref. (1). Conditional on non-singleton variants, the average length of the tree, denoted by $\bar{T}$, is equal to $n(n+1)/2$ and the average coalescence time of lineage $i$ and lineage $j$ can be described as

$$\bar{t}(i,j) = \begin{cases} \max(i,j)\,, \text{ if } i \neq j\,, \\ 0\,, \text{ otherwise.} \end{cases}$$

To find the eigenvectors and eigenvalues of $\mathbb{E}[\mathbf{K}]$, we study the spectrum of matrix $\bar{T}\mathbb{E}[\mathbf{K}]/n^2$. We look for an eigenvector $\mathbf{u}$ and the corresponding eigenvalue $\mu$ such that

$$\frac{\bar{T}}{n^2}\mathbb{E}[\mathbf{K}]\mathbf{u} = \mu\mathbf{u}\,.$$

We set $\lambda = n^2\mu/\bar{T}$, so that we have $\mathbb{E}[\mathbf{K}]\mathbf{u} = \lambda\mathbf{u}$. The eigenvalue $\lambda$ can be approximated as $\lambda \approx 2\mu$. To solve the eigenvalue problem, we seek solutions of the form $\mathbf{u}(i) = u(i/n)$, $i = 1, \ldots, n$, where $u(x)$ is a continuous function determined by

$$\left(\frac{\bar{t}(i)}{n} - \frac{\bar{t}}{n}\right)\times\frac{1}{n}\sum_{j=1}^{n} u(j/n) + \frac{1}{n}\sum_{j=1}^{n} u(j/n)\frac{\bar{t}(j)}{n} - \frac{1}{n}\sum_{\substack{j=1\\ i\neq j}}^{n} \max(i/n, j/n)u(j/n) = \mu u(i/n). \tag{3}$$

The average coalescence time of lineage $i$ and a random lineage divided by $n$ can be approximated as

$$\frac{\bar{t}(i)}{n} = \frac{1}{n}\sum_{\substack{k=1\\ k\neq i}}^{n} \max(i/n, k/n) = \frac{1}{n}\sum_{k=1}^{n} \max(i/n, k/n) - \frac{i}{n^2} \approx \frac{x^2+1}{2} - \frac{x}{n},$$

and the average coalescence time of two random lineages divided by $n$ can be approximated by

$$\frac{\bar{t}}{n} = \frac{1}{n^2}\sum_{k=1}^{n}\sum_{\ell=1}^{n} \max(\ell/n, k/n) - \frac{1}{n^2}\sum_{k=1}^{n} k/n \approx \frac{2}{3} - \frac{1}{2n}\,.$$

The discrete eigenvalue problem (3) becomes a continuous equation for large $n$

$$\int_0^1 u(y)\left[\frac{x^2+y^2}{2} + \frac{1}{3} - \frac{x+y}{n} + \frac{1}{2n} - \max(x,y)\right]dy = \mu u(x) - \frac{x}{n}u(x). \tag{4}$$

We define a function $k$ such that

$$k(x,y) = \frac{x^2+y^2}{2} + \frac{1}{3} - \frac{x+y}{n} + \frac{1}{2n} - \max(x,y).$$

By direct integration, we have

$$\int_0^1 k(x,y)dx = -y/n.$$

Integrating both sides of the equation (4), it comes

$$\int_0^1 \int_0^1 u(y)k(x,y)dydx = \int_0^1 u(x)\left(\mu - \frac{x}{n}\right)dx,$$

exchanging order of integrals, we have

$$\int_0^1 u(y)dy = 0. \tag{5}$$

Therefore, the problem becomes

$$\frac{1}{2}\int_0^1 u(y)\left(y^2 - \frac{2y}{n}\right)dy - \int_0^1 u(y)\max(x,y)dy = \mu u(x) - \frac{x}{n}u(x). \tag{6}$$

A first differentiation of equation (6) gives

$$-\int_0^x u(y)dy = \mu u'(x) - u'(x)\frac{x}{n} - u(x)\frac{1}{n}, \tag{7}$$

and a second differentiation leads us to

$$u''(x)\left(\mu - \frac{x}{n}\right) - u'(x)\frac{2}{n} + u(x) = 0. \tag{8}$$

We look for the solution of the equation (8), we set $a = n\mu - x$ and define the function $f(a)$ such that $f(a) = f(n\mu - x) = u(x)$, then equation (8) becomes

$$f''(a)a + 2f'(a) + nf(a) = 0.$$

We assume $x < \mu n$ for all $x \in [0,1]$, so that $a$ is always positive. We set $h(a) = \sqrt{a}f(a)$. The successive derivatives of the function $f$ are given by

$$\begin{cases} f(a) = h(a)a^{-1/2} \\ f'(a) = h'(a)a^{-1/2} - \frac{1}{2}h(a)a^{-3/2} \\ f''(a) = h''(a)a^{-1/2} - \frac{1}{2}h'(a)a^{-3/2} - \frac{1}{2}h'(a)a^{-3/2} + \frac{3}{4}h(a)a^{-5/2} \end{cases}$$

Using the derivatives of the function $f$, we get

$$h''(a)a^{1/2} + h'(a)a^{-1/2} + h(a)\left[na^{-1/2} - \frac{1}{4}a^{-3/2}\right] = 0.$$

Multiplying by $a$, we have

$$h''(a)a^{3/2} + h'(a)a^{1/2} + h(a)\left[na^{1/2} - \frac{1}{4}a^{-1/2}\right] = 0.$$

After simplification, we have

$$h''(a)\frac{4u^2}{4\sqrt{a}} + h'(a)\frac{4u}{4\sqrt{a}} + h(a)\left[\frac{4na}{4\sqrt{a}} - \frac{1}{4\sqrt{a}}\right] = 0.$$

Multiplying by the denominators, we get

$$h''(a)a^2 + h'(a)a + h(a)\left[na - 1/4\right] = 0. \tag{9}$$

We set $a = v^2/4n$ and define a function $\nu(v)$ such that $\nu(v) = \nu(2\sqrt{n}\sqrt{a}) = h(a)$. The successive derivatives of the function $\nu(v)$ are given by

$$\begin{cases} \partial_u \nu(v) = \partial_u \nu(\sqrt{a}2\sqrt{n}) = 2\sqrt{n}\frac{1}{2\sqrt{a}}\partial_v \nu(v) = \frac{\sqrt{n}}{\sqrt{a}}\partial_v \nu(v) \\ \partial_u^2 \nu(v) = \partial_u\left[\frac{\sqrt{n}}{\sqrt{a}}\partial_v \nu(v)\right] = -\frac{1}{2}\sqrt{n}a^{-3/2}\partial_v \nu(v) + \frac{\sqrt{n}}{\sqrt{a}}2\sqrt{n}\frac{1}{2\sqrt{a}}\partial_v^2 \nu(v) \end{cases} \tag{10}$$

Then, we have

$$\underbrace{\nu''(v)4na - 2\sqrt{na}\nu'(v)}_{h''(a)(2a)^2} + \nu'(v)\underbrace{4\sqrt{na}}_{2ah'(a)} + \nu(v)\left[4na - 1\right] = 0.$$

Replacing $a$ by $v^2/2n$, the above equation is transformed into the following Bessel differential equation

$$\nu''(v)v^2 + v\nu'(v) + \nu(v)\left[v^2 - 1\right] = 0.$$

The solution of the differential equation has the following form

$$\nu(v) = AJ_1(v) + BY_1(v),$$

where $J_1$ is the Bessel function of the first kind and $Y_1$ is the Bessel function of the second kind, and $A$ and $B$ are constant values determined by boundary conditions described later.

Substituting $v$ with $x$ and $\nu$ with $u$, we have

$$u(x) = A\frac{J_1(2\sqrt{n}\sqrt{\mu n - x})}{\sqrt{\mu n - x}} + B\frac{Y_1(2\sqrt{n}\sqrt{\mu n - x})}{\sqrt{\mu n - x}}. \tag{11}$$

Using equation (7), we have two boundary conditions

$$\begin{cases} \mu u'(0) = \frac{u(0)}{n} \approx 0 \\ \mu u'(1) = \frac{u'(1)}{n} + \frac{u(1)}{n} \approx 0 \end{cases} \tag{12}$$

The derivative of the function $u(x)$ is equal to

$$\begin{aligned} u'(x) = &-\frac{A}{\mu n - x}\left(\sqrt{n}J_0(2\sqrt{n}\sqrt{\mu n - x}) - \frac{1}{\sqrt{\mu n - x}}J_1(2\sqrt{n}\sqrt{\mu n - x})\right) \\ &- \frac{B}{\mu n - x}\left(\sqrt{n}Y_0(2\sqrt{n}\sqrt{\mu n - x}) - \frac{1}{\sqrt{\mu n - x}}Y_1(2\sqrt{n}\sqrt{\mu n - x})\right). \end{aligned} \tag{13}$$

To simplify equation (13), we use the following recurrence identities

$$J_n(x) = \frac{2(n+1)}{x}J_{n+1}(x) - J_{n+2}(x), \quad Y_n(x) = \frac{2(n+1)}{x}Y_{n+1}(x) - Y_{n+2}(x), \quad \text{for } n \in \mathbb{N}. \tag{14}$$

Then, equation (13) becomes

$$u'(x) = A\frac{\sqrt{n}}{\mu n - x}J_2(2\sqrt{n}\sqrt{\mu n - x}) + B\frac{\sqrt{n}}{\mu n - x}Y_2(2\sqrt{n}\sqrt{\mu n - x}). \tag{15}$$

Substituting this into equations (12) gives the following two conditions

$$\begin{cases} AJ_2(2\sqrt{n}\sqrt{\mu n}) + BY_2(2\sqrt{n}\sqrt{\mu n}) = 0 \\ AJ_2(2\sqrt{n}\sqrt{\mu n - 1}) + BY_2(2\sqrt{n}\sqrt{\mu n - 1}) = 0 \end{cases}$$

The eigenvalues $\mu$ greater than $1/n$ are defined by the zeros of the function

$$\mu \mapsto J_2(2\sqrt{n}\sqrt{\mu n})Y_2(2\sqrt{n}\sqrt{\mu n - 1}) - Y_2(2\sqrt{n}\sqrt{\mu n})J_2(2\sqrt{n}\sqrt{\mu n - 1}).$$

It follows that the eigenvalues $\lambda$ greater than $2/n$ are defined by the zeros of the function

$$\lambda \mapsto J_2(\sqrt{2\lambda}n)Y_2(\sqrt{2(\lambda n - 2)n}) - Y_2(\sqrt{2\lambda}n)J_2(\sqrt{2(\lambda n - 2)n}).$$

#### 2 Normalizing constant for the first PC eigenvectors

In this section we prove the following formula of the normalizing constant $A_k$ for the eigenvectors of the unfiltered data analysis.

$$\begin{aligned}\frac{1}{A_k^2} = &\Big(J_1(\sqrt{2(\lambda_k n - 2)n}) - \rho_k Y_1(\sqrt{2(\lambda_k n - 2)n})\Big)^2 \left[1 + \frac{2}{n(\lambda_k n - 2)}\right] \\ &- \Big(J_1(\sqrt{2\lambda_k}n) - \rho_k Y_1(\sqrt{2\lambda_k}n)\Big)^2 \left[1 + \frac{2}{n^2\lambda_k}\right], \quad (16)\end{aligned}$$

where $\rho_k = J_2(\sqrt{2\lambda_k}n)/Y_2(\sqrt{2\lambda_k}n)$.

According to (1), the $k$th eigenvector have the following form, for $k \in 1, \ldots, n$,

$$u_k(x) = A_k \left( \frac{J_1(2\sqrt{n}\sqrt{\mu_k n - x})}{\sqrt{\mu_k n - x}} - \frac{J_2(2\sqrt{n}\sqrt{\mu_k n})}{Y_2(2\sqrt{n}\sqrt{\mu_k n})} \frac{Y_1(2\sqrt{n}\sqrt{\mu_k n - x})}{\sqrt{\mu_k n - x}} \right),$$

To normalize the $k$th eigenvector we define $A_k$ such that

$$\frac{1}{A_k^2} = \int_0^1 u_k^2(y)dy,$$

where

$$\begin{aligned}
u_k^2(x) &= \frac{J_1(2\sqrt{n}\sqrt{\mu_k n - x})^2}{\mu_k n - x} + \frac{Y_1(2\sqrt{n}\sqrt{\mu_k n - x})^2}{\mu_k n - x}\frac{J_2(2\sqrt{n}\sqrt{\mu_k n})^2}{Y_2(2\sqrt{n}\sqrt{\mu_k n})^2} \\
&- 2\frac{J_1(2\sqrt{n}\sqrt{\mu_k n - x})Y_1(2\sqrt{n}\sqrt{\mu_k n - x})}{\mu_k n - x}\frac{J_2(2\sqrt{n}\sqrt{\mu_k n})}{Y_2(2\sqrt{n}\sqrt{\mu_k n})} \\
&= u_k^{(1)}(x) + u_k^{(2)}(x) + u_k^{(3)}(x).
\end{aligned}$$

Using the linearity of the integral, we have

$$\int_0^1 u_k(x)^2 dx = \int_0^1 u_k^{(1)}(x)dx + \int_0^1 u_k^{(2)}(x)dx + \int_0^1 u_k^{(3)}(x)dx = (1) + (2) + (3).$$

We compute the following integrals

$$\int_0^1 \frac{J_1(2\sqrt{n}\sqrt{\mu_k n - x})^2}{\mu_k n - x}dx, \quad \int_0^1 \frac{Y_1(2\sqrt{n}\sqrt{\mu_k n - x})^2}{\mu_k n - x}dx, \quad \int_0^1 \frac{J_1(2\sqrt{n}\sqrt{\mu_k n - x})Y_1(2\sqrt{n}\sqrt{\mu_k n - x})}{\mu_k n - x}dx \tag{17}$$

Setting $u = \mu_k n - x$, the integrals (17) becomes

$$\int_{\mu_k n-1}^{\mu_k n} \frac{J_1(2\sqrt{n}\sqrt{u})^2}{u}du, \quad \int_{\mu_k n-1}^{\mu_k n} \frac{Y_1(2\sqrt{n}\sqrt{u})^2}{u}du, \quad \int_{\mu_k n-1}^{\mu_k n} \frac{J_1(2\sqrt{n}\sqrt{u})Y_1(2\sqrt{n}\sqrt{u})}{u}du.$$

Setting $x = 2\sqrt{n}\sqrt{u}$, it follows that the above terms are equal to

$$2\int_{2\sqrt{n}\sqrt{\mu_k n-1}}^{2\sqrt{n}\sqrt{\mu_k n}} \frac{J_1(x)^2}{x}dx, \quad 2\int_{2\sqrt{n}\sqrt{\mu_k n-1}}^{2\sqrt{n}\sqrt{\mu_k n}} \frac{Y_1(x)^2}{x}dx, \quad 2\int_{2\sqrt{n}\sqrt{\mu_k n-1}}^{2\sqrt{n}\sqrt{\mu_k n}} \frac{J_1(x)Y_1(x)}{x}dx.$$

For all real numbers $a$ and $b$ such that $b > a > 0$, we have the following identities

$$\begin{cases} \int_a^b \frac{J_1(y)^2}{y}\, dy = \frac{1}{2}\left(J_0(a)^2 + J_1(a)^2 - J_0(b)^2 - J_1(b)^2\right) \\ \int_a^b \frac{Y_1(y)^2}{y}\, dy = \frac{1}{2}\left(Y_0(a)^2 + Y_1(a)^2 - Y_0(b)^2 - Y_1(b)^2\right) \\ \int_a^b \frac{J_1(y)Y_1(y)}{y}\, dy = \frac{1}{2}\left(J_0(a)Y_0(a) + J_1(a)Y_1(a) - J_0(b)Y_0(b) - J_1(b)Y_1(b)\right) \end{cases}$$

Thus, we have

$$(1) = J_0\left(2\sqrt{n}\sqrt{\mu_k n - 1}\right)^2 + J_1\left(2\sqrt{n}\sqrt{\mu_k n - 1}\right)^2 - J_0\left(2\sqrt{n}\sqrt{\mu_k n}\right)^2 - J_1\left(2\sqrt{n}\sqrt{\mu_k n}\right)^2,$$

$$\begin{aligned}(2) = \frac{J_2(2\sqrt{n}\sqrt{\mu_k n})^2}{Y_2(2\sqrt{n}\sqrt{\mu_k n})^2}\Bigg( & Y_0\left(2\sqrt{n}\sqrt{\mu_k n - 1}\right)^2 + Y_1\left(2\sqrt{n}\sqrt{\mu_k n - 1}\right)^2 \\ & - Y_0\left(2\sqrt{n}\sqrt{\mu_k n}\right)^2 - Y_1\left(2\sqrt{n}\sqrt{\mu_k n}\right)^2\Bigg),\end{aligned}$$

$$\begin{aligned}(3) = -2\frac{J_2(2\sqrt{n}\sqrt{\mu_k n})}{Y_2(2\sqrt{n}\sqrt{\mu_k n})}\Bigg( & J_0(2\sqrt{n}\sqrt{\mu_k n - 1})Y_0(2\sqrt{n}\sqrt{\mu_k n - 1}) \\ & + J_1(2\sqrt{n}\sqrt{\mu_k n - 1})J_1(2\sqrt{n}\sqrt{\mu_k n - 1})\Bigg) \\ + 2\frac{J_2(2\sqrt{n}\sqrt{\mu_k n})}{Y_2(2\sqrt{n}\sqrt{\mu_k n})} & \left[J_0(2\sqrt{n}\sqrt{\mu_k n})Y_0(2\sqrt{n}\sqrt{\mu_k n}) - J_1(2\sqrt{n}\sqrt{\mu_k n})Y_1(2\sqrt{n}\sqrt{\mu_k n})\right],\end{aligned}$$

We set $a = 2\sqrt{n}\sqrt{\mu_k n - 1}$, $b = 2\sqrt{n}\sqrt{\mu_k n}$ and $\rho = J_2(b)/Y_2(b)$. Therefore, we have

$$(1)+(2)+(3) = \left(J_0(a) - \rho Y_0(a)\right)^2 + \left(J_1(a) - \rho Y_1(a)\right)^2 - \left(J_0(b) - \rho Y_0(b)\right)^2 - \left(J_1(b) - \rho Y_1(b)\right)^2.$$

Then, we have

$$(1) + (2) + (3) = \left(J_1(a) - \rho Y_1(a)\right)^2\left[1 + \frac{4}{a^2}\right] - \left(J_1(b) - \rho Y_1(b)\right)^2\left[1 + \frac{4}{b^2}\right].$$

Replacing $a$, $b$ by their values defined above, this concludes the proof of equation (16)
